# Daughterless, the *Drosophila* orthologue of TCF4, is required for associative learning and maintenance of synaptic proteome

**DOI:** 10.1101/792796

**Authors:** Laura Tamberg, Mariliis Jaago, Kristi Säälik, Anastassia Shubina, Carl Sander Kiir, Alex Sirp, Tõnis Timmusk, Mari Palgi

## Abstract

Mammalian Transcription Factor 4 (TCF4) has been linked to schizophrenia and intellectual disabilities like Pitt-Hopkins syndrome (PTHS). Here we show that similarly to mammalian TCF4, fruit fly orthologue Daughterless (Da) is expressed in the *Drosophila* brain structures associated with learning and memory, the mushroom bodies. Furthermore, silencing of *da* in mushroom body neurons impairs appetitive associative learning of the larvae and leads to decreased levels of the synaptic proteins Synapsin (Syn) and discs large 1 (dlg1) suggesting the involvement of Da in memory formation. Here we demonstrate that *Syn* and *dlg1* are direct target genes of Da in adult *Drosophila* heads, since Da binds to the regulatory regions of these genes and the modulation of Da levels alter the levels of *Syn* and *dlg1* mRNA. Silencing of *da* also affects negative geotaxis of the adult flies suggesting the impairment of locomotor function. Overall, our findings suggest that Da regulates *Drosophila* larval memory and adult negative geotaxis possibly via its synaptic target genes *Syn* and *dlg1*. These behavioural phenotypes can be further used as a PTHS model to screen for therapeutics.

**Summary statement:** Human TCF4, a bHLH transcription factor, is associated with intellectual disability and schizophrenia. Here we propose a *Drosophila* model for human disease studies using TCF4 orthologue in fruit fly, Daughterless.

## Introduction

Transcription Factor 4 (TCF4, also known as ITF2, E2-2, SEF2 etc.) belongs to the family of class I basic helix-loop-helix (bHLH) transcription factors, also called E-proteins (Murre et al., 1994). The E-proteins bind to the DNA Ephrussi box (E-box) sequence CANNTG as homodimers or heterodimers with class II bHLH transcription factors (Cabrera and Alonso, 1991). TCF4, Transcription factor 4, should be distinguished from T cell factor 4, also called TCF4 with official name TCF7L2, interacting with β-catenin and participating in WNT signalling pathway. TCF4 is essential for a range of neurodevelopmental processes including early spontaneous neuronal activity, cell survival, cell cycle regulation, neuronal migration and differentiation, synaptic plasticity, and memory formation (Chen et al., 2016; Crux et al., 2018; Forrest et al., 2013; Hill et al., 2017; Jung et al., 2018; Kennedy et al., 2016; Kepa et al., 2017; Li et al., 2019; Page et al., 2018; Thaxton et al., 2018). Genes involved in pathways including nervous system development, synaptic function and axon development are TCF4 targets (Forrest et al., 2018; Xia et al., 2018). Furthermore, TCF4 regulates the expression of ion channels Na_V_1.8 and K_V_7.1 (Ekins et al., 2019; Rannals et al., 2016). Recent insights into the mechanisms of activation of TCF4 show that TCF4-dependent transcription in primary neurons is induced by neuronal activity via soluble adenylyl cyclase and protein kinase A (PKA) signalling (Sepp et al., 2017). In addition to nervous system, TCF4 has been shown to function in immune system in plasmacytoid dendritic cells development (Cisse et al., 2008; Grajkowska et al., 2017).

Deficits in TCF4 function are associated with several human diseases. TCF4 haploinsufficiency causes Pitt-Hopkins syndrome (PTHS; OMIM #610954) (Amiel et al., 2007; Brockschmidt et al., 2007; Zweier et al., 2007). Patients with PTHS have severe intellectual disability, developmental delay, intermittent hyperventilation periods followed by apnea, and display distinct craniofacial features, reviewed in international consensus statement (Zollino et al., 2019). Currently there is no treatment for PTHS, but dissecting the functional consequences triggered by mutated TCF4 alleles could reveal attractive avenues for curative therapies for this disorder (reviewed in Rannals and Maher, 2017). Large scale genome wide association studies revealed SNPs in *TCF4* among the top risk loci for schizophrenia (SCZ) (Talkowski et al., 2012). Consistently, *TCF4* is involved in SCZ endophenotypes like neurocognition and sensorimotor gating (Lennertz et al., 2011a; Lennertz et al., 2011b; Quednow et al., 2011). Furthermore, many genes that are mutated in SCZ, autism spectrum disorder and intellectual disability patients are TCF4 target genes (Forrest et al., 2018). How deficits in TCF4 function translate into neurodevelopmental impairments and whether TCF4 plays an essential role in the mature nervous system is poorly understood.

We have previously demonstrated that TCF4 function can be modelled in *Drosophila melanogaster* using its orthologue and the sole E-protein in the fruit fly – Daughterless (Da). PTHS-associated mutations introduced to Da lead to similar consequences in the fruit fly as do the same mutations in TCF4 *in vitro* (Sepp et al., 2012; Tamberg et al., 2015). Furthermore, human TCF4 is capable of rescuing the lack of Da in the development of *Drosophila* embryonic nervous system (Tamberg et al., 2015). Da is involved in various developmental processes including sex determination, neurogenesis, myogenesis, oogenesis, intestinal stem cell maintenance, and development of the eye, trachea and salivary gland (Bardin et al., 2010; Bhattacharya and Baker, 2011; Brown et al., 1996; Castanon et al., 2001; Caudy et al., 1988; Cline, 1978; Cummings and Cronmiller, 1994; King-Jones et al., 1999; Massari and Murre, 2000; Smith et al., 2002; Wong et al., 2008). In the developing nervous system the role of Da is well established during neuronal specification as an obligatory heterodimerization partner for proneural class II bHLH transcription factors (Cabrera and Alonso, 1991; Powell et al., 2008). However, the functional role of Da following neurogenesis and nervous system maturation remains unknown.

Here we set out to characterize the expression of Da in the nervous system. To this end we created *Drosophila* lines where Da protein was endogenously tagged with either 3xFLAG or sfGFP epitope tags. We show that Da is broadly expressed in the larval CNS including mushroom body, the memory and learning centre of insects. To test whether Da is involved in learning and memory formation in the fruit fly we used appetitive associative learning paradigm in larvae (Michels et al., 2017). In this assay, reduced levels of Da in the mushroom body resulted in impaired learning and memory formation. Furthermore, silencing of *da* also resulted in decreased level of synaptic proteins Synapsin (Syn) and discs large 1 (dlg1). Therefore, we suggest that the knockdown of *da* in mushroom body neurons combined with appetitive associative learning paradigm is further applicable to screen for potential therapeutics for the treatment of PTHS as well putative genetic interactors of Da and by proxy, TCF4. We also demonstrate that Da binds to several areas in the *dlg1* gene and to *Syn* promoter region in adult *Drosophila* heads and that overexpression of *da* increases *Syn* and *dlg1* mRNA levels in the adult heads. Therefore, we have shown for the first time that Da is required to sustain elements of the synaptic proteome in a mature nervous system positing a post-developmental function for Da and possibly TCF4.

## Results

### Da is expressed in all developmental stages of the fruit fly

While the expression of Da protein has been studied in fruit fly embryos, ovaries, larval optic lobes and imaginal discs using various anti-Da antibodies (Andrade-Zapata and Baonza, 2014; Bhattacharya and Baker, 2011; Bhattacharya and Baker, 2012; Brown et al., 1996; Cronmiller and Cummings, 1993; Li and Baker, 2018; Tanaka-Matakatsu et al., 2014; Yasugi et al., 2014), its expression during adulthood remains largely uncharacterized. Therefore, we first aimed to study Da expression throughout the development of the fruit fly using immunoblot analysis. Since there are no commercial antibodies available that recognize Da we used the CRISPR/Cas9 system to create transgenic flies where Da is N-terminally tagged with 3xFLAG epitope. The resulting 3xFLAG-*da* line is maintained as homozygotes indicating that the tagged Da protein is functional as both *da* null mutations and *da* ubiquitous overexpression lead to embryonic lethality (Caudy et al., 1988; Giebel et al., 1997). We then characterized Da expression throughout development and in adult *Drosophila* heads of the 3xFLAG-*da* line by performing immunoblot analysis with anti-FLAG antibodies. During development, we compared Da expression from embryonic to late pupal stages (Figure 1A and 1C). In adults, we analysed the Da levels from the heads of one, four and seven day old males and females (Figure 1B and 1D). We found Da to be expressed throughout development, with the highest levels detected in the 3rd instar larvae and early pupae while the lowest levels were observed in the 2nd instar larvae and late pupae (Figure 1C). During adulthood, Da expression was highest in the heads of one day old females and decreased thereafter in both males and females (Figure 1D). Also, one day old females had significantly higher Da expression in the head than one day old males (Figure 1D).

**Figure 1.**
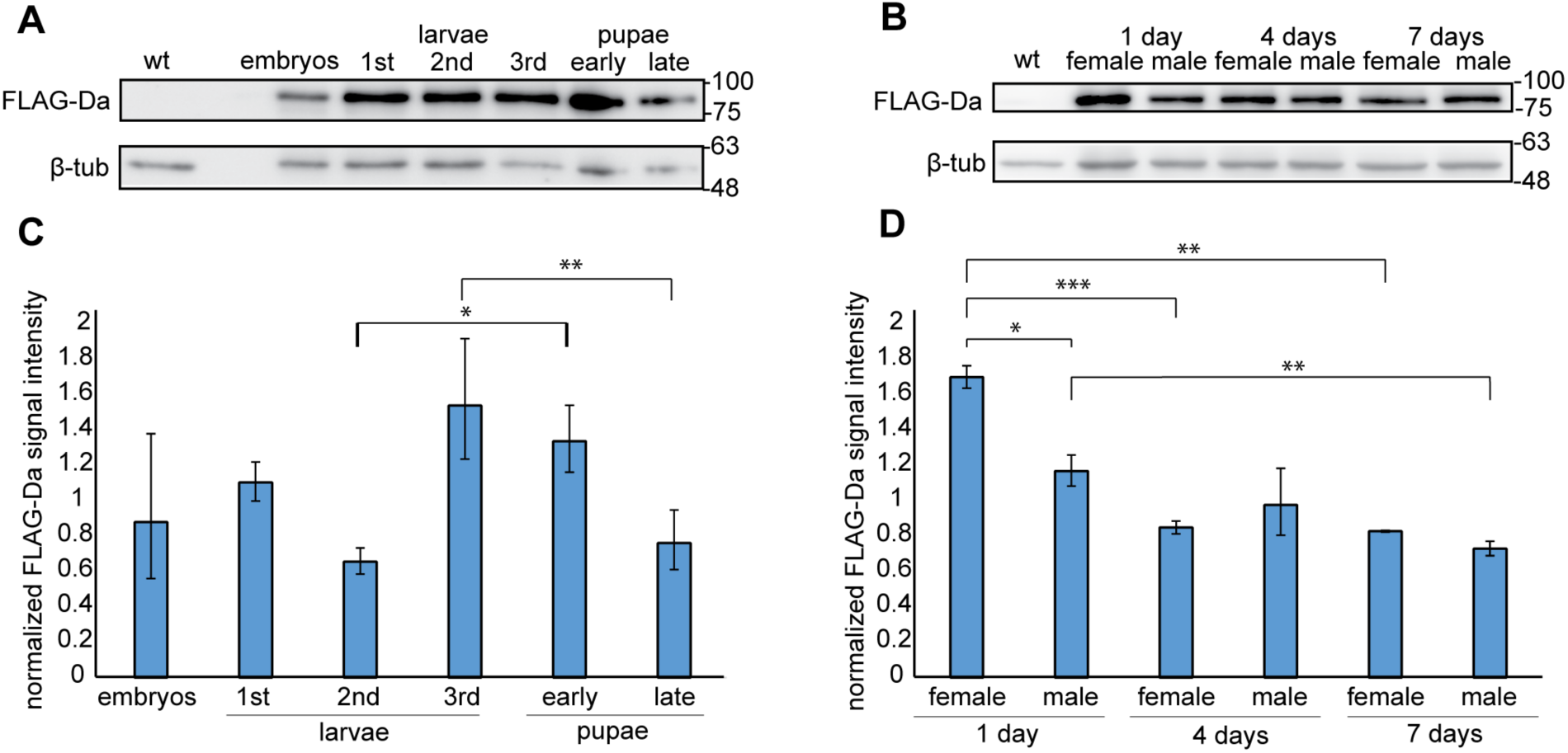
Da is expressed in all developmental stages of the fruit fly. A and B - 3xFLAG-Da fusion protein is expressed throughout the fruit fly development. Western blot analysis was carried out using anti-FLAG antibody, w^1118^ wild type - wt serves as negative control, numbers on the right side indicate molecular weights of proteins in kDa. C and D - results of densitometric analysis of Western blot, 3xFLAG-Da signals were normalized using β-tubulin signals. The mean results from three independent Western blots are shown. Error bars show standard errors. Statistical significance is shown with asterisks between the groups connected with lines. *P<0.05, **P<0.01, ***P<0.001, Student t-test.

### 3xFLAG-Da retains transactivational capability of Da in HEK293 cells

In addition to 3xFLAG-*da* we also created sfGFP-*da* flies, where *da* is tagged with superfolder green fluorescent protein (sfGFP) in the same N-terminal position. To determine whether N-terminal tagging of Da proteins influences their transactivation capability we used the luciferase reporter system where the expression of the *luciferase* gene is controlled by E-boxes with a minimal promoter. For this, we cloned the 3xFLAG-tagged or sfGFP-tagged *da* from the genomes of the tagged lines into mammalian expression vector pcDNA3.1 and overexpressed these constructs in HEK293 cells. Luciferase reporter assay showed that compared to wild type (wt) Da the transactivational capability of 3xFLAG-Da was unchanged (Figure 2A). In contrast, the transactivational capability of sfGFP-Da was significantly reduced (Figure 2A). Both *3xFLAG-da* and *sfGFP*-*da* constructs were expressed at equal levels as revealed by Western blot analysis (Figure 2B). This suggests that the 3xFLAG tag does not interfere with the transcriptional activity of Da.

**Figure 2.**
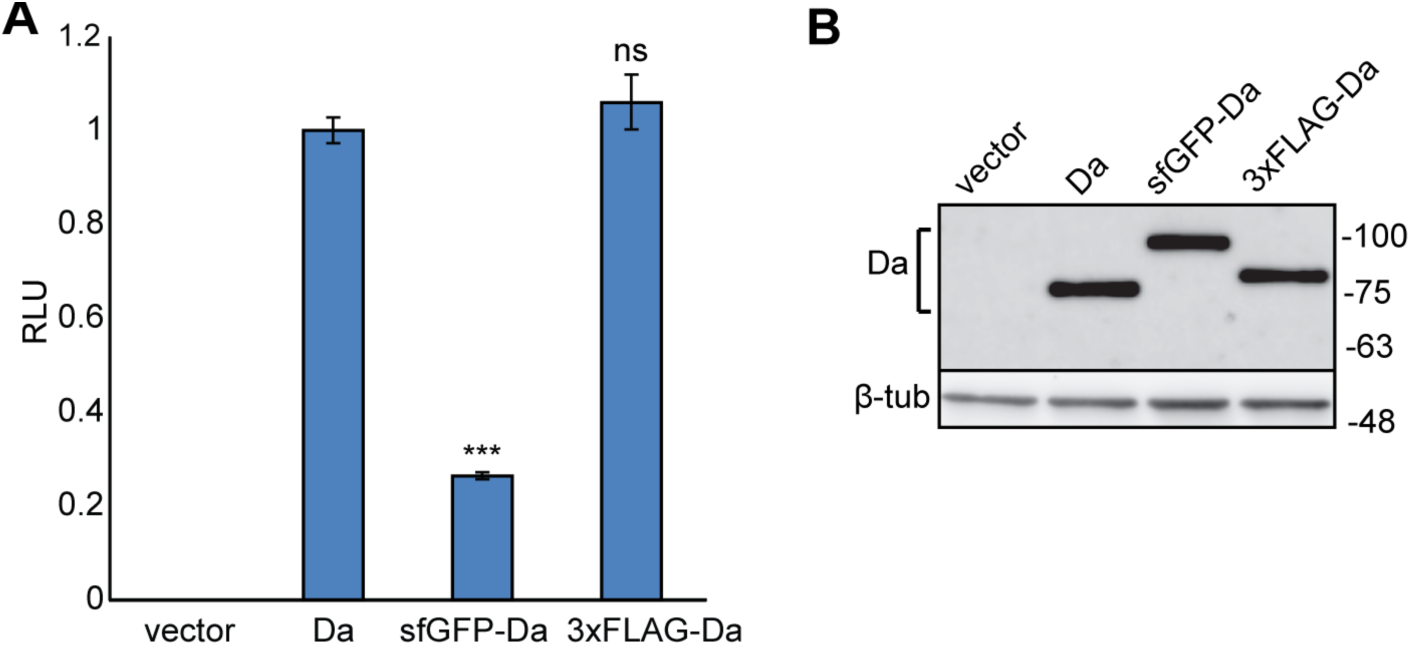
Transactivational capability of Da is unaffected by N-terminal 3xFLAG tag, but is reduced by sfGFP tag. A - HEK293 cells were co-transfected with constructs encoding wild type *da*, tagged *da*, or empty vector, firefly luciferase construct carrying 12 µE5 boxes with a minimal promoter, and *Renilla* luciferase construct without E-boxes for normalization. Luciferase activities were measured and data are presented as fold induced levels compared to the signals obtained from cells transfected with wild type *da* encoding construct. The mean results from six independent transfection experiments performed in duplicates are shown. Error bars show standard errors. Statistical significance is shown with asterisks relative to wt Da expressing cells ***P<0.001, ns - not significant, Student t-test; RLU - relative luciferase unit. B - Western blot from transfected HEK293 cells using anti-Da antibody dam109-10. Wild type Da, sfGFP- and 3xFLAG-tagged Da are all expressed at equal levels. Numbers on the right side indicate molecular weight of proteins in kDa.

### Da is expressed in the brain structures associated with learning and memory

Next we used the 3xFLAG-*da* line to characterize expression of Da in the third instar larval brain. Da was expressed throughout the larval central nervous system (CNS), with stronger expression detected in the optic lobes (Figure 3A’’ - 3C’’). Since mutations in or deletion of one of the *TCF4* alleles lead to PTHS in humans and one of the hallmarks of PTHS is severe learning disability, and since *TCF4* is expressed in the adult mammalian hippocampus (Jung et al., 2018), we aimed to determine whether TCF4 homolog Da is expressed in the mushroom body, the brain structure of insects responsible for learning and memory. For this, we deployed the UAS-Gal4 binary expression system (Brand and Perrimon, 1993) by combining the 3xFLAG-*da* line with different driver lines with expression in the mushroom body. Resulting lines with 3xFLAG-*da* and Gal4 were then combined with nuclear targeted UAS-nls-GFP. The GMR12B08-Gal4 line directed expression of Gal4 under the control of the single intron of *da* in most regions of the brain, including the mushroom body (Figure 3A). The two other drivers, 30Y-Gal4 and 201Y-Gal4, were both mushroom body-specific (Figure 3B and 3C). We observed that the expression of 3xFLAG-Da and GMR12B08>nlsGFP overlapped in many areas of the third instar larval brain (Figure 3A’), including the mushroom body (Figure 3B’ and 3C’). In contrast, with 30Y-Gal4 and 201Y-Gal4, the mushroom body-specific driver lines, 3xFLAG-Da showed partial co-expression in cells contributing to the third instar mushroom body. Thus, Da is expressed broadly in the CNS of 3rd instar larvae including mushroom body.

**Figure 3.**
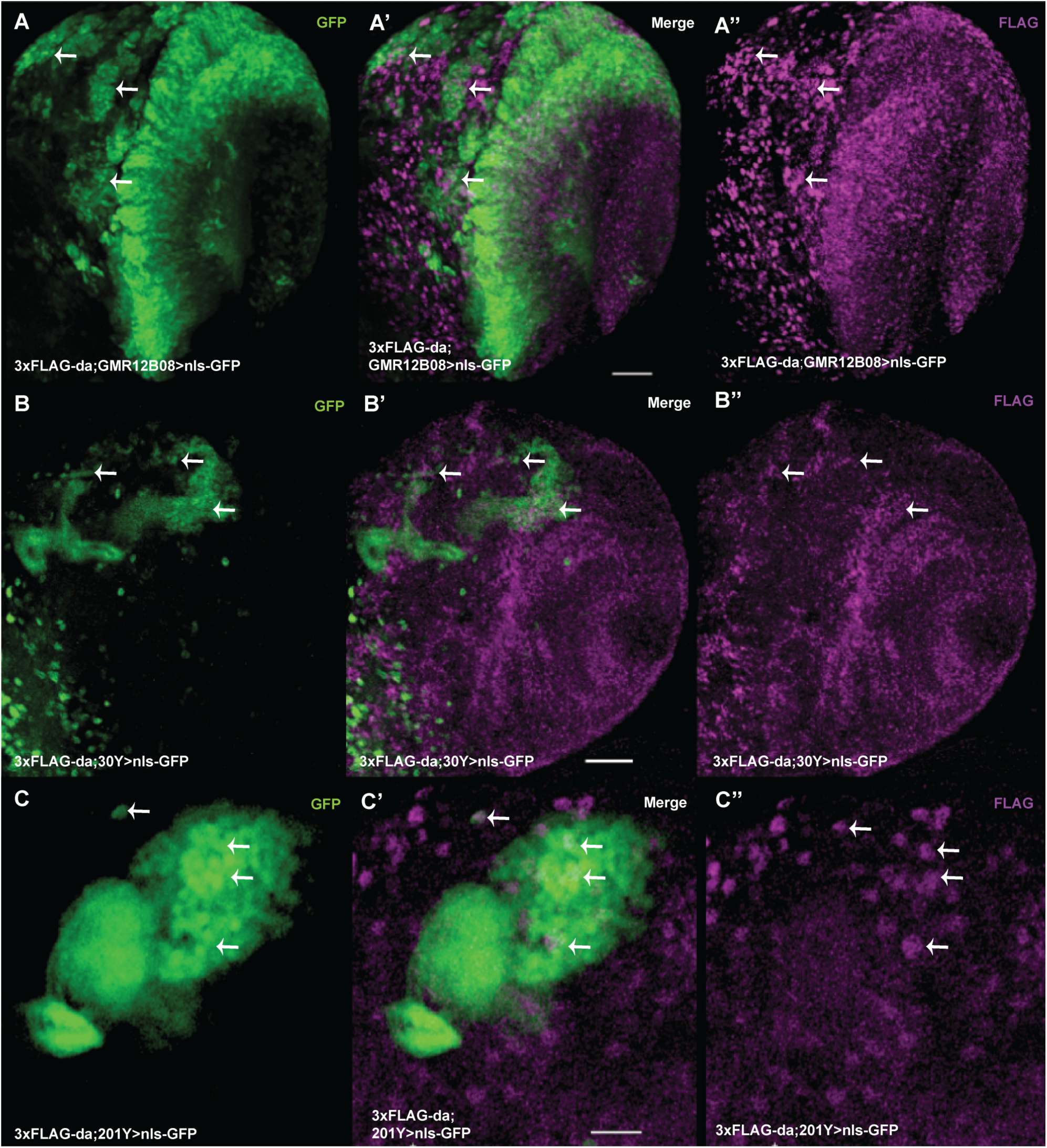
In third instar larval brain Da is expressed widely including the mushroom body. Single slices of laser confocal microscopy presented. Da is coexpressed with GMR12B08-Gal4 in many areas of the larval brain lobe (A-A’’). Da is expressed in the mushroom body, which is marked by nlsGFP driven by 30Y-Gal4 (B-B’’) and by 201Y-Gal4 (C-C’’). A, B, and C - nlsGFP expression shows driver expression pattern. A’’, B’’ and C’’ - expression of 3xFLAG-Da. Some sites of coexpression of nlsGFP and 3xFLAG-Da are shown by arrows. Scale bars on A’ and B’ represent 30 µm and on C’ 20 µm.

### Silencing of *da* in the CNS leads to impaired memory of the larvae

Heterozygous mutations in *TCF4*, the orthologue of *da*, lead to the PTHS syndrome, characterized by intellectual disability. This fact, and the observation that Da is expressed in the mushroom body implies that it might be involved in learning and memory in flies. To test this, we decided to take advantage of the ease of assaying appetitive associative learning and memory in the *Drosophila* larvae (Michels et al., 2017). However, this assay showed that learning ability is not impaired in *da* heterozygous mutants (Figure 4A), which could be due to *da* upregulation by autoregulation (Smith and Cronmiller, 2001). Since sfGFP-Da showed diminished transactivation capability *in vitro* (see above), we also tested homozygous sfGFP-*da* larvae, and found no impairment of learning ability (Figure 4A). Thus, we next investigated whether knockdown of *da* with concurrent enhancement by Dicer-2 (Dcr2) expression (Dietzl et al., 2007) in the *Drosophila* CNS could impact memory and learning ability. To silence *da* in the CNS, we used several CNS-specific Gal4 lines. We found that appetitive associative learning was impaired in flies when using three drivers - GMR12B08-Gal4 (Figure 4B) and mushroom body specific lines 30Y-Gal4 (Figure 4C) and 201Y-Gal4 (Figure 4D). For control experiments, we used both the UAS-*da*^*RNAi*^ line and the UAS-*Dcr2* driven by the CNS-specific Gal4 line. All of the transgenes (UAS-*Dcr2*, UAS-*da*^*RNAi*^ and the driver Gal4) were homozygous for lowered learning ability in the case of GMR12B08-Gal4 and 201Y-Gal4 (Figure 4B and 4D). In the case of UAS-*Dcr2*;*3xFLAG-da*,UAS-*da*^*RNAi*^;GMR12B08-Gal4 line (where *3xFLAG-da*, UAS-*da*^*RNAi*^ and GMR12B08-Gal4 were all in a homozygous state), Da levels in the larval brains were reduced by approximately 25% and 35% when compared to UAS-Dcr2;3xFLAG-*da*;GMR12B08-Gal4 and 3xFLAG*-da* larval brains, respectively (Suppl. figure 1A and 1B). We were unable to get homozygotes using mushroom body specific driver 30Y-Gal4, possibly because knockdown of *da* with this driver in homozygous state is too strong and causes lethality. The learning ability of heterozygous larvae where *da* was silenced with the 30Y-Gal4 driver was notably but non-significantly impaired (Figure 4C). Larvae with impaired learning were also tested for their ability to taste and smell. All three lines showed preference towards fructose (Suppl. fig 2A) and amyl-acetate (Suppl. fig 2B) which indicates that the larvae can sense taste and smell and that their locomotor activity is not affected. Expressing only Dcr2 under neuron-specific *elav-Gal4*, as a control for *da* silencing using the same driver, as such already caused learning impairment (data not shown), so learning ability of *elav-Gal4>Dcr2;da*^*RNAi*^ larvae was nonsignificantly different from the control larvae *elav-Gal4>Dcr2*. Knockdown of *da* with glia-specific *repo-Gal4* did not alter learning and memory (data not shown). This suggests that for normal larval appetitive associative memory appropriate Da levels are needed in the brain structures specified by GMR12B08-Gal4, 30Y-Gal4 and 201Y-Gal4.

**Figure 4.**
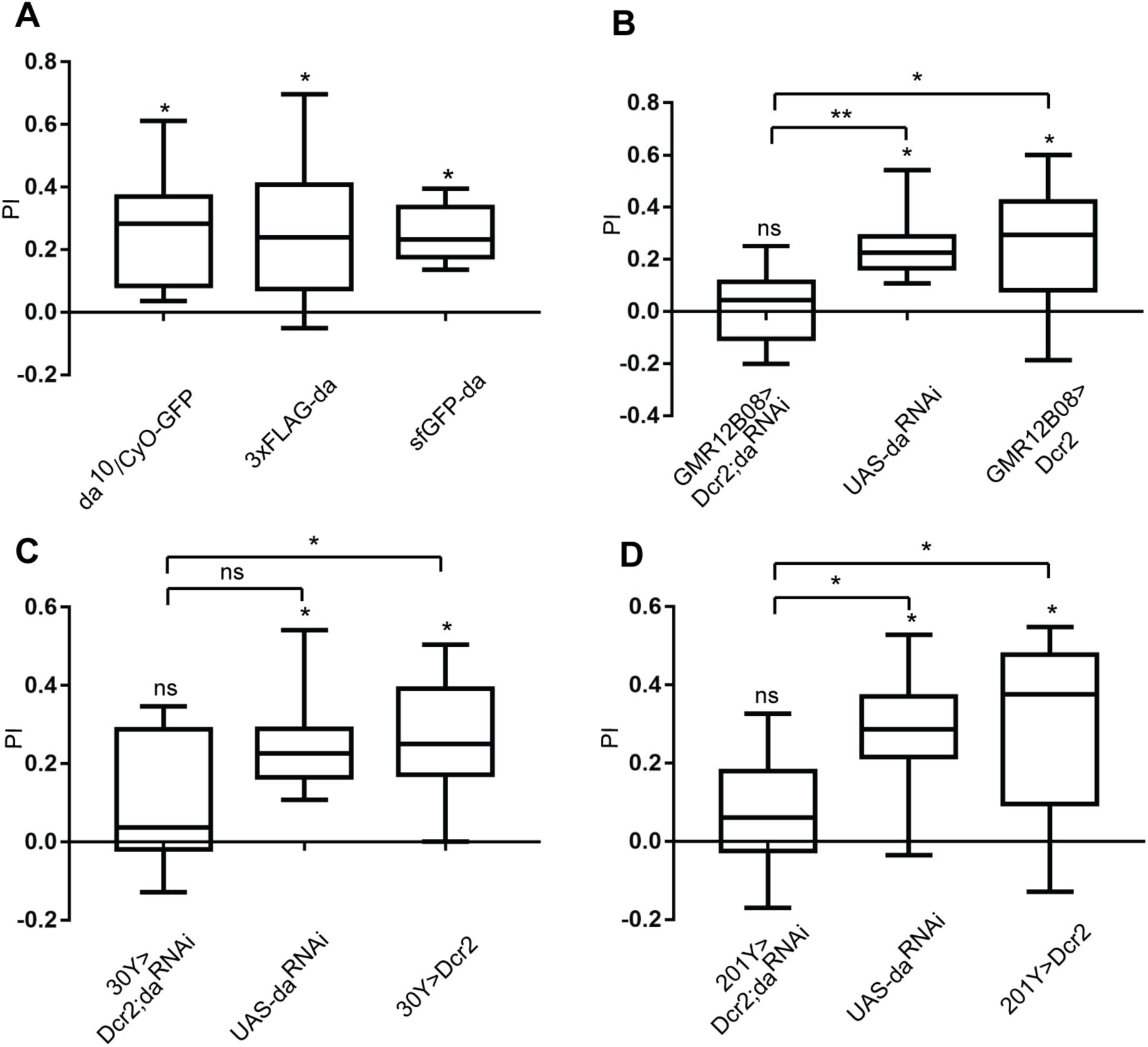
Knockdown of *da* in the mushroom body leads to impaired olfactory learning of larvae. A - Heterozygous *da* mutation does not cause reduction of appetitive associative learning, and both FLAG-*da* and GFP-*da* larvae show olfactory learning. B - Appetitive associative learning was impaired when *da* was silenced using GMR12B08-Gal4, C - 30Y-Gal4 and D - 201Y-Gal4. UAS-*Dcr2*, UAS-*da*^*RNAi*^ and Gal4 - all were homozygous when driven by GMR12B08-Gal4 (B) or 201Y-Gal4 (D). In the case of 30Y-Gal4 homozygotes never emerged and the effect on learning was smaller (C). PI-s (performance index) are visualized using box-whisker plots, which show the median, the 10% - 90% quantiles, and the 25% - 75% quantiles. For statistical analysis one-sample sign test and Mann-Whitney U test with Bonferroni correction were used. * p<0.025, ** p<0.005.

### Reduced level of Da in the larval CNS leads to decreased expression of synaptic proteins Synapsin and discs large 1

To investigate the putative mechanisms underlying learning and memory deficits in larvae with lowered levels of Da in the nervous system we used the driver line GMR12B08-Gal4 for silencing *da* since it had the broadest expression. For this we compared the expression levels of several known synaptic proteins in the 3rd instar larval brains under both *da* knockdown and overexpression conditions using the GMR12B08-Gal4 line (Figure 5). We quantified the expression levels of presynaptic protein bruchpilot (brp) (Figure 5A), postsynaptic protein discs large 1 (dlg1) (Figure 5B), presynaptic Synapsin (Syn) (Figure 5C) which is important for learning and memory (Michels et al., 2005), and pan-neuronally expressed neuronal specific splicing factor embryonic lethal abnormal vision (elav) (Figure 5D). We found that the levels of both dlg1 and Syn were reduced in 3rd instar larval brains with lower levels of Da (Figure 5B and 5C respectively). On the other hand, Da overexpression did not result in increased levels of these proteins. The levels of elav and brp were not significantly changed by knockdown or overexpression of *da* (Figure 5A and 5D, respectively). The finding that elav levels were not affected by Da suggests that reducing Da levels does not affect the number of neurons and the observed learning impairment might rather stem from lowered expression levels of synaptic proteins or, alternatively, from reduced number of synapses.

**Figure 5.**
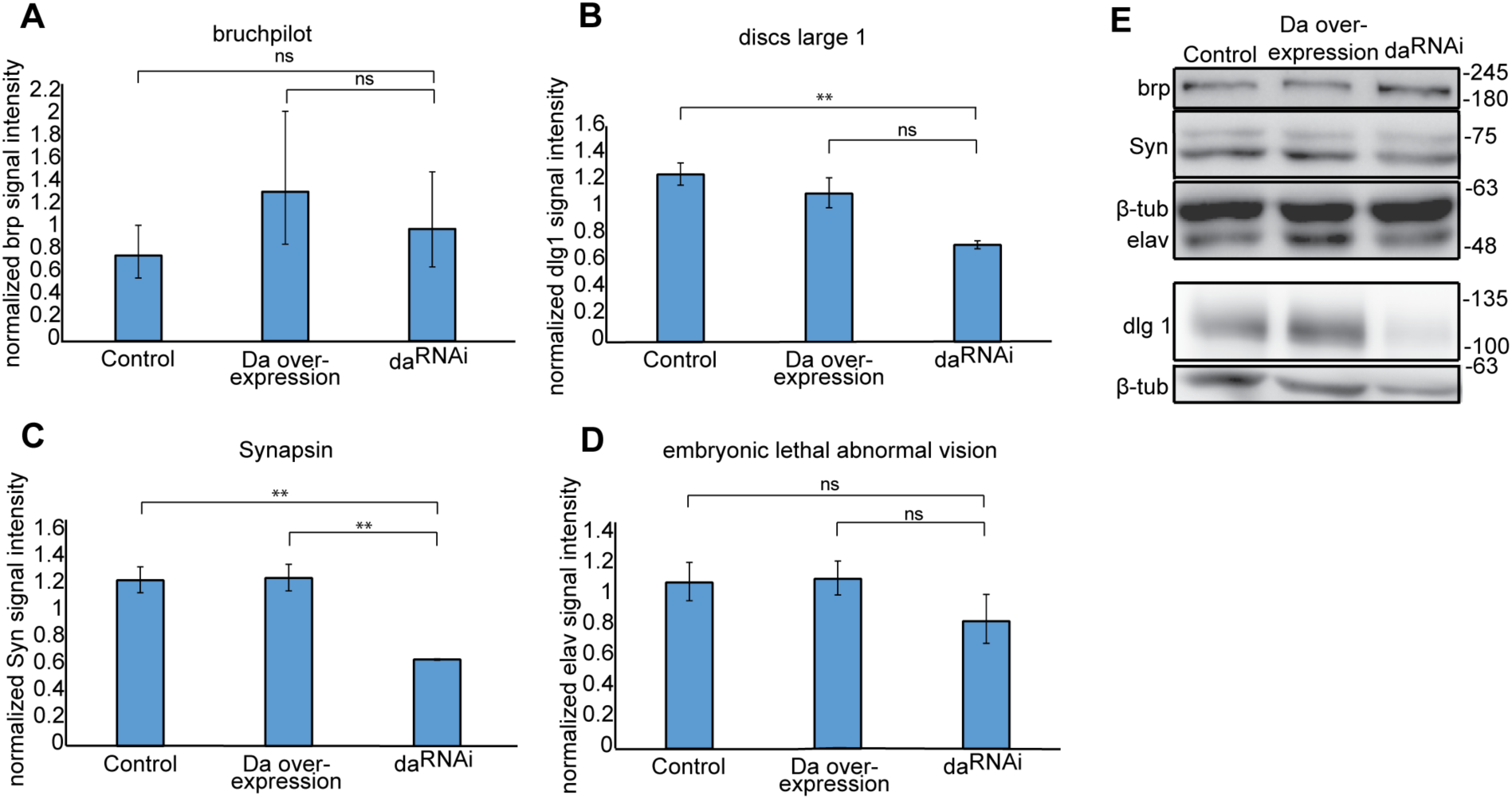
Silencing of Da lowers expression levels of Synapsin and discs large 1. Western blot was carried out using larval brains where *da* was silenced with GMR12B08-Gal4. A-D - results of densitometric analysis of Western blot, protein signals were normalized using β-tubulin signals. The mean results from four independent Western blots are shown. Error bars show standard errors. Statistical significance is shown with asterisks between the groups connected with lines. *P<0.05, **P<0.01, ***P<0.001, Sudent t-test. Overexpression of Da does not alter bruchpilot, discs large 1, Synapsin or embryonic lethal abnormal vision levels (A-D). discs large 1 and Synapsin expression levels are lower when Da is silenced (B and C respectively). Silencing of Da does not change bruchpilot and embryonic lethal abnormal vision expression levels (A and D respectively). E – Representative Western blot using 3rd instar larval brains. Numbers indicate molecular weight of proteins in kDa. Control - GMR12B08>Dcr2 larval brains, Da overexpression - GMR12B08>Da larval brains, da^RNAi^ - GMR12B08>Dcr2,da^RNAi^ larval brains.

### Larval appetitive associative learning and memory test can be used for screening drugs for PTHS treatment

Our finding showing that larval appetitive associative learning and memory becomes impaired upon *da* silencing indicates that these fly lines could be used for modelling certain aspects of PTHS in *Drosophila* and testing potential drug candidates. For instance, various drugs or drug candidates could be tested for their capacity to rescue this behavioural impairment. Previously, it has been shown in our group that resveratrol improves transactivational capability of TCF4 in primary neuronal cultures (unpublished data). Also histone deacetylase inhibitor suberoylanilide hydroxamic acid (SAHA) has been shown to rescue memory impairment in the mouse model of PTHS (Kennedy et al., 2016). We therefore decided to test these two substances in the appetitive associative learning experiments. We observed that while the *da* knockdown larvae fed with 400 µM resveratrol or 2 µM SAHA showed increased associative memory, the rescue of the learning deficit was nonsignificant when compared to the controls (Figure 6A and 6B). Nevertheless, the impaired memory and learning of *Drosophila* larvae with specific *da* knockdown can be further used for screening pharmaceuticals for potential treatment of PTHS.

**Figure 6.**
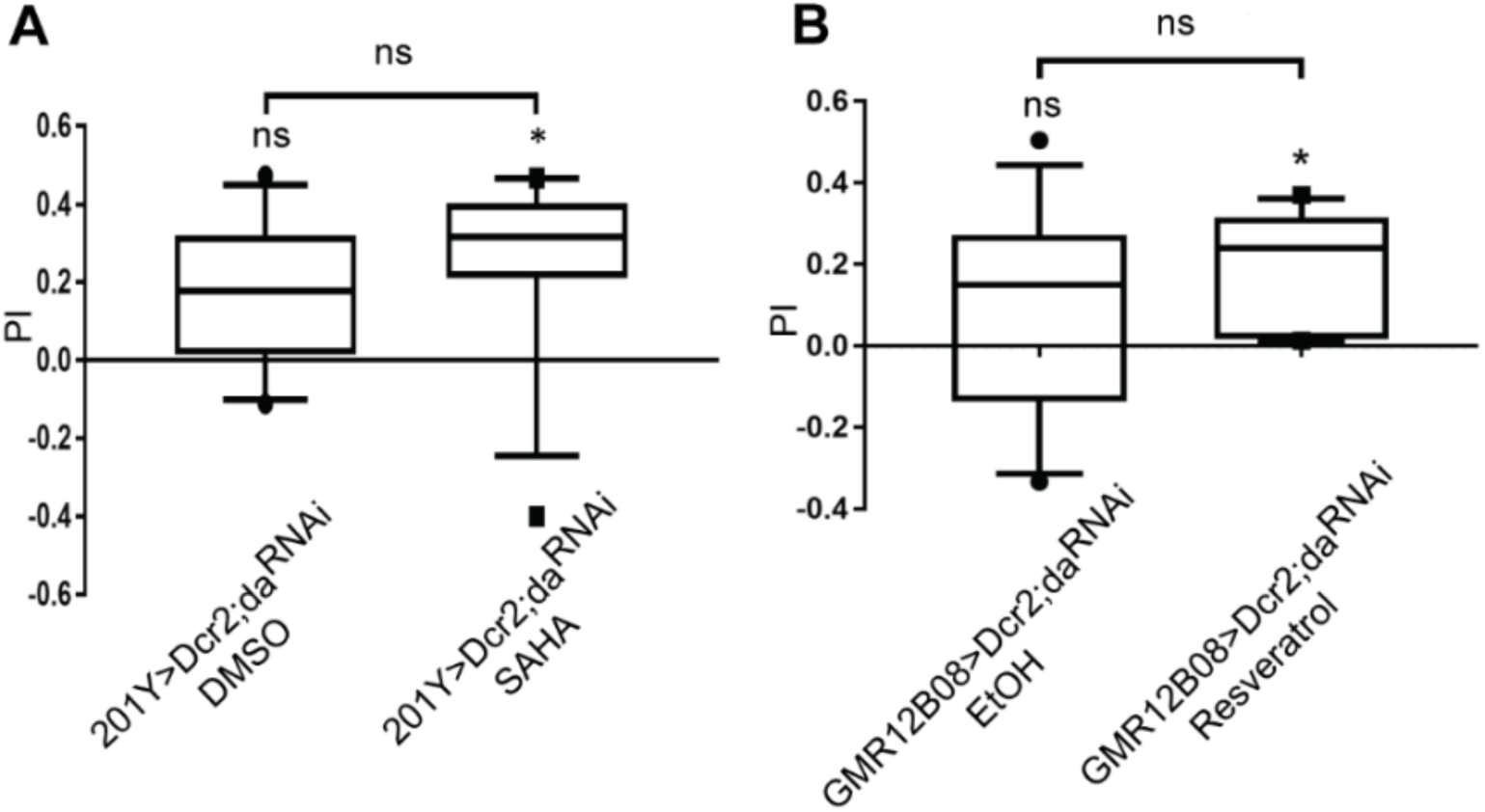
Resveratrol and SAHA have moderate positive effects on rescuing the impaired learning phenotype resulting from decreased levels of Da. A and B - Adding 400 µM resveratrol or 2 µM SAHA to the larval growth media improves appetitive associative memory. PI-s are visualized using box-whisker plots, which show the median, the 10% - 90% quantiles, and the 25% - 75% quantiles. For statistical analysis one-sample sign test and Mann-Whitney U test with Bonferroni correction were used. Median PI (performance index) of these larvae are significantly different from zero so these larvae show associative memory although the PI-s are not significantly different compared to the controls.

### Suppressing Da using 30Y-Gal4 leads to impaired negative geotaxis of adult flies

Negative geotaxis has been successfully used to evaluate climbing ability indicative of motor dysfunction in the *Drosophila* model for Angelman syndrome that has similar symptoms to PTHS (Wu et al., 2008). Thus, we next used this assay to evaluate locomotion in adult flies where *da* knockdown had been achieved by the same drivers as used for the larval learning test. We found that negative geotaxis was unchanged in homozygotes where *da* knockdown had been achieved by the broad neuronal driver GMR12B08-Gal4 or mushroom body specific driver 201Y-Gal4 (Figure 7A or 7C respectively). Interestingly, both female and male heterozygotes whose *da* was silenced by the 30Y-Gal4 driver had severely impaired negative geotaxis (Figure 7B). Next we visualized Da expression in the adult heads using 3xFLAG-*da* line and confirmed its coexpression with 30Y-Gal4. Da was expressed widely in the adult *Drosophila* brain including the central brain and optic lobes (Figure 7D-D’’), and coexpressed with 30Y-Gal4 in many cells in the mushroom body (Figure 7E-E’’). 30Y-Gal4 is expressed in all mushroom body αβ, α’β’ and γ lobes but 201Y-Gal4 is not expressed in α’β’ Kenyon cells (Aso et al., 2009). It has previously been shown that when mushroom body α’β’ Kenyon cells are activated then negative geotaxis is decreased (Sun et al., 2018). Silencing *da* by 201Y-Gal4 or GMR12B08-Gal4 did not cause decreased negative geotaxis probably because these drivers are not expressed in brain areas responsible for this behaviour, for example mushroom body α’β’ lobes. We also made an attempt to rescue this phenotype by administrating resveratrol or SAHA either to larvae during development or to adult flies in the food substrate, but we could not get an improvement (data not shown).

**Figure 7.**
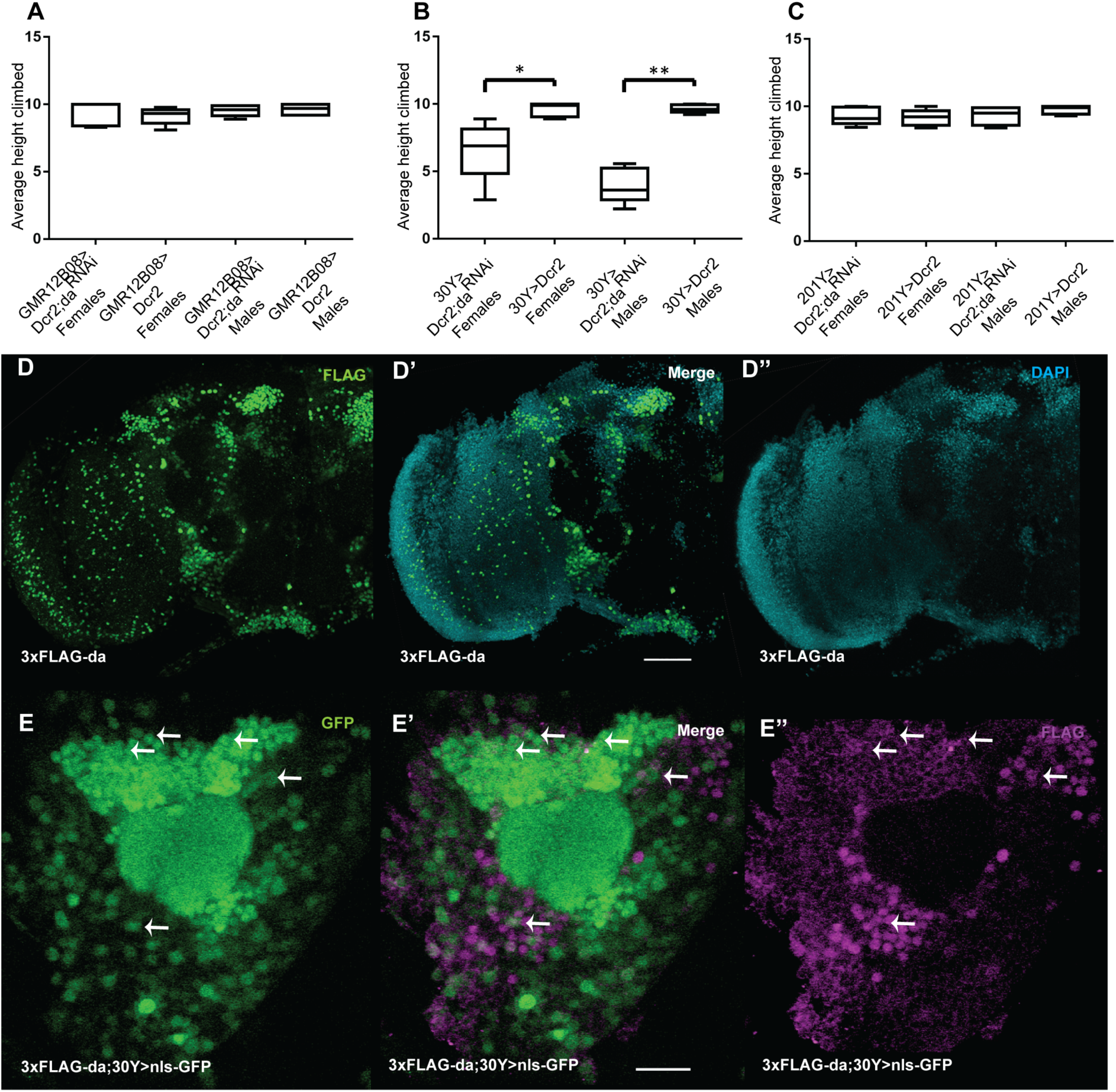
Silencing of *da* with 30Y-Gal4 impairs negative geotaxis in adult flies. Negative geotaxis was not affected when Da was suppressed using GMR12B08-Gal4 (A) or 201Y-Gal4 (C). The climbing height of the flies was significantly lower when Da was silenced using 30Y-Gal4 (B). Average climbing heights are visualized using box-whisker plots, which show the median, the 10% - 90% quantiles, and the 25% - 75% quantiles. For statistical significance pairwise U-tests were used. * p<0.05, ** p<0.01. D-D’’ – Da is expressed widely in the adult *Drosophila* brain including the central brain and optic lobes. 3xFLAG-Da is green on D and D’ and nuclei are visualized using DAPI on D’ and D’’. Scale bar represents 50 µm (D’). E-E’’ – Da coexpresses with 30Y-Gal4 in adult *Drosophila* mushroom body. 30Y-Gal4 expression is visualized with nlsGFP (E and E’) and 3xFLAG-Da is magenta (E’ and E’’). Some parts of coexpression is shown by arrows. Scale bar represents 15 µm (E’).

### *Synapsin* and *discs large 1* are Da target genes

Since Syn and dlg1 levels were reduced in third instar larval brains when *da* was silenced and it has been shown that Da binds to both *Syn* and *dlg1* gene locus in embryonic stage 4-5 (MacArthur et al., 2009) we sought to investigate whether Da binds to these areas in adult heads too. For this we used chromatin immunoprecipitation assay (ChIP) in 3xFLAG-*da* adult heads using anti-FLAG antibodies. As a control we used *white*^*1118*^ fly line with no FLAG tag. For quantitative PCR (qPCR) from immunoprecipitated chromatin we designed primers to amplify *Syn* and *dlg1* gene areas containing E-boxes where Da binds in early embryos (MacArthur et al., 2009). In addition to previously shown Da binding site in *Syn* gene we also tested Da binding to *Syn* promoter region (Figure 8A). For *dlg1* we designed four primer pairs, since Da has been shown to bind four areas in that gene (Figure 8B) (MacArthur et al., 2009). As a negative control we used primers for *achaete* (Andrade-Zapata and Baonza, 2014) since achaete is a proneural protein essential for neuronal development and should not be expressed in adult heads. As a positive control we used *peptidylglycine alpha-hydroxylating monooxygenase (PHM)* gene first intron where Da binds as a heterodimer with dimmed to activate transcription (Park et al., 2008). qPCR from immunoprecipitated chromatin using *Syn* primers resulted in enrichment of *Syn* promoter area (primer pair SynI) while previously reported Da binding site was not enriched in adult heads (primer pair SynII) (Figure 8C). All *dlg1* primers resulted in enrichment of previously reported Da binding areas (Figure 8C). This means that Da does not bind to the locus at the 3’ end of *Syn* gene but binds to *Syn* promoter and all four *dlg1* gene areas that we selected in the adult heads. To validate *Syn* and *dlg1* as Da target genes we carried out RT-qPCR analysis in adult *Drosophila* heads under *da* silencing and overexpression conditions. Although upon *da* silencing using *elav-Gal4* only *Syn* mRNA levels were decreased (Figure 8D), *da* overexpression using *elav-Gal4* increased mRNA levels of both *Syn* and *dlg1* (Figure 8E). This indicates that both *Syn* and *dlg1* are direct targets of Da in the *Drosophila* nervous system.

**Figure 8.**
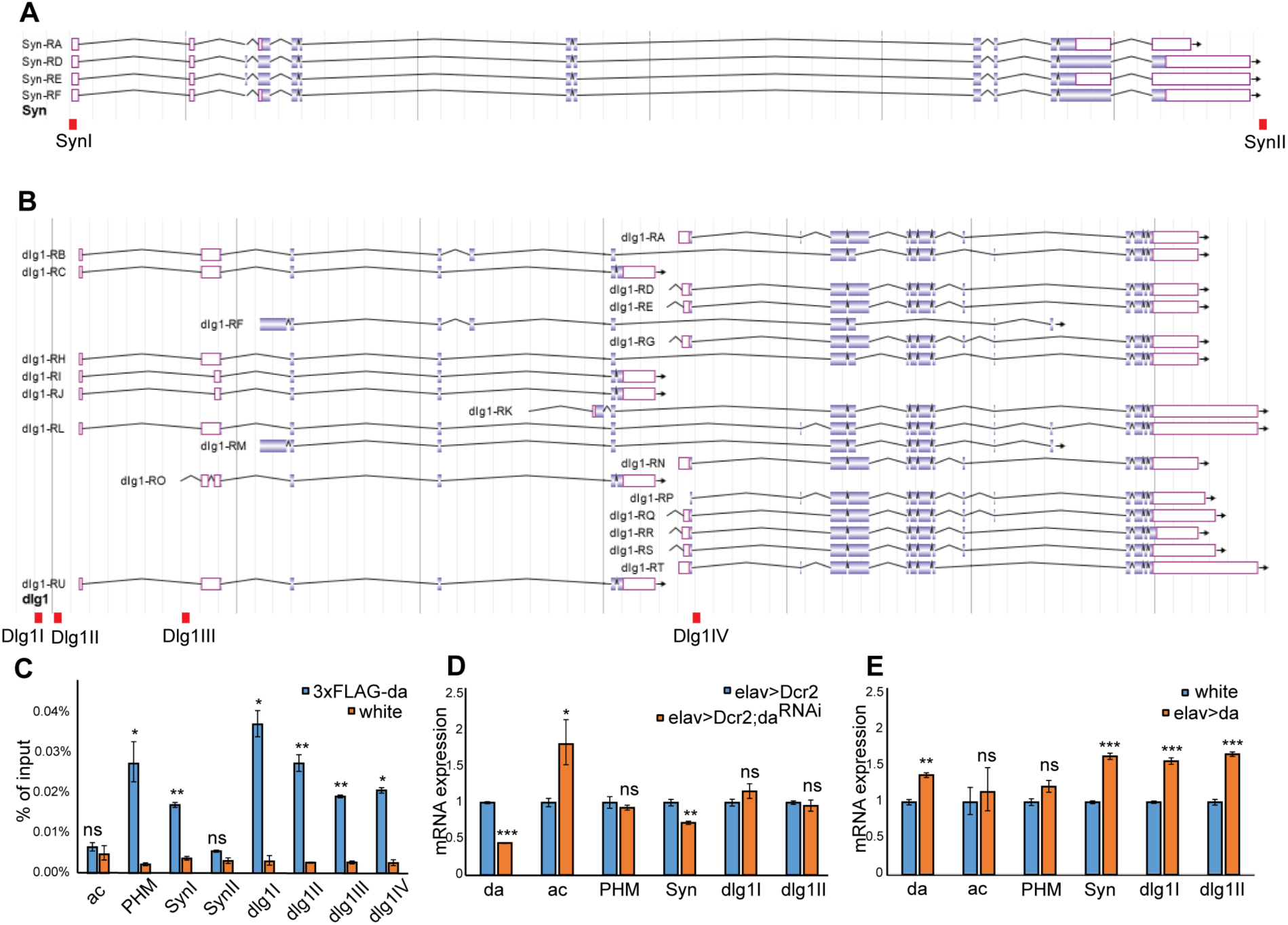
Da directly regulates *Synapsin* and *discs large 1* in adult fly heads. A – JBrowse view of *Syn* gene, four annotated transcripts are shown. B – JBrowse view of *dlg1* gene, 21 annotated transcripts are shown. Red boxes indicate areas, where primer pairs Syn1, SynII, Dlg1I, Dlg1II, Dlg1III and Dlg1IV amplify DNA. C - qPCR results from chromatin immunoprecipitation experiment from 3xFLAG-*da* and *white*^*1118*^ wild type adult heads using anti-FLAG antibody. ac – *achaete* gene locus as a negative control; PHM - *peptidylglycine alpha-hydroxylating monooxygenase* gene first intron as a positive control; SynI – promoter region of *Synapsin*, SynII – 3’ end of *Synapsin* gene locus; dlg1I, dlg1II, dlg1III, dlg1IV – *discs large 1* gene locus. D, E – RT-qPCR results showing the effects of *da* silencing (D) or overexpression (E) using elav-Gal4 on *da, ac, PHM, Syn* and *dlg1* mRNA levels. *da* silencing reduces *da* and *Syn*, and increases *ac* mRNA levels (D). *da* overexpression increases *da, Syn* and *dlg1* mRNA levels (E). C, D, E - Results from three biological replicates are shown, error bars indicate standard errors. *P<0.05, **P<0.01, ***P<0.001, Student t-test.

## Discussion

Here we characterized the expression of Da in the *Drosophila melanogaster* larval and adult brain. Da was expressed in many areas of the brain including the mushroom body, which is the centre for learning and memory in the fruit fly and carries out a role that is comparable to the mammalian hippocampus. Orthologue of *da, TCF4* is expressed not only in the adult mammalian hippocampus but also in cortical and subcortical structures (Jung et al., 2018).

We created N-terminally tagged 3xFLAG-*da* and sfGFP-*da* fly strains. Both strains are homozygous viable and fertile, indicating that the overall functionality of Da *in vivo* is not altered by the molecular tag. However, in luciferase reporter assay in mammalian HEK293 cells the sfGFP tag reduced transcription activation capability of Da. E-proteins activate transcription preferably as heterodimers with class II bHLH proteins, but can also act as homodimers (Cabrera and Alonso, 1991). In mammalian HEK293 cells Da has to activate transcription as a homodimer, since there are no heterodimerization partners like class II bHLH proteins expressed (Sepp et al., 2012). This suggests that sfGFP tag could interfere with Da function as a homodimer in the luciferase assay but not as a heterodimer *in vivo*. We also compared appetitive associative learning ability of 3xFLAG-*da* and sfGFP-*da* larvae and both of the lines had no learning impairment in this assay. This provides additional evidence that the 3xFLAG tag does not affect Da function and sfGFP tag reduces its transactivational capability probably by interfering with Da homodimer function.

Since Da is expressed in larval brain structures associated with learning and memory, and PTHS is caused by heterozygous mutations in *TCF4* we tested learning ability of *da* heterozygous mutant larvae. These larvae had no memory impairment, which could be due to *da* upregulation by autoregulation (Smith and Cronmiller, 2001). Learning and memory of larvae was impaired when *da* was knocked down in the mushroom body. Da mammalian orthologue TCF4 is also associated with learning and memory, since when *TCF4* is downregulated in mouse hippocampus, pathways associated with neuronal plasticity are dysregulated (Kennedy et al., 2016) and silencing of *TCF4* in human pluripotent stem cell-derived neurons results in downregulated signalling pathways important for learning and memory (Hennig et al., 2017). In *TCF4* conditional knock-out mice the neurons in the cortex and hippocampus have reduced numbers of dendritic spines, which also suggests that synaptic plasticity is altered (Crux et al., 2018). In multiple PTHS mouse models spatial learning is defective probably through hippocampal NMDA receptor hyperfunction (Thaxton et al., 2018). Furthermore, many genes that code for synaptic proteins and have been linked to autism, intellectual disability, or psychiatric diseases are direct targets of TCF4 (Forrest et al., 2018; Hennig et al., 2017).

Here we show that when *da* is silenced using driver with broad expression in the *Drosophila* larval brain, expression levels of synaptic proteins disc large 1 (dlg1) and Synapsin (Syn) are downregulated. dlg1 is a member of the membrane-associated guanylate kinase (MAGUK) protein family. Several vertebrate homologs of dlg1 have been shown to be important for learning and memory. Discs Large MAGUK Scaffold Protein 3 (DLG3), also called Synapse-Associated Protein 102 (SAP-102) knock-out mice have spatial learning deficit (Cuthbert et al., 2007) and in human *DLG3* mutations which cause dysfunctional NMDA receptor signalling have been associated with X-linked mental retardation (Tarpey et al., 2004; Zanni et al., 2010). We also found that Da is a direct regulator of *dlg1*, since in the adult *Drosophila* heads Da binds to multiple areas in *dlg1* gene and *dlg1* expression is upregulated when *da* is overexpressed. Gene coding for Discs Large MAGUK Scaffold Protein 2 (DLG2) also called Postsynaptic Density Protein 93 (PSD-93) which is a homolog of *Drosophila dlg1*, is a direct target of TCF4 (Hennig et al., 2017), which indicates that Da and TCF4 share at least some common mechanisms in regulating learning and memory.

Synapsins are presynaptic phosphoproteins that regulate synaptic output (reviewed in Diegelmann et al., 2013). There are three genes coding for vertebrate Synapsins but only one *Syn* gene in *Drosophila* (Klagges et al., 1996). Using knockout experiments in mice it has been shown that Synapsins are involved in learning and memory (Gitler et al., 2004; Silva et al., 1996) and *SYN1* has been implicated in human neurological diseases such as learning difficulties and epilepsy (Garcia et al., 2004). Likewise, *Drosophila syn*^*97*^ mutant larvae have impaired appetitive associative learning (Michels et al., 2005). The fact that memory of *syn*^*97*^ larvae can be rescued by expressing Syn in the mushroom bodies (Michels et al., 2011) is consistent with our findings that lower Da levels affect Syn expression levels and that appropriate Da levels in mushroom body are required for proper memory formation. This means that Da and Syn-dependent memory trace is formed in the mushroom bodies. Syn-dependent memory is likely formed by its phosphorylation by Protein kinase A (PKA) (Michels et al., 2011). When Syn is phosphorylated at its PKA/CamK I/IV (Protein kinase A/ Ca^2+^/calmodulin-dependent protein kinase I/IV) sites its affinity for actin is reduced and synaptic vesicles from the reserve pool can be exocytosed (reviewed in Benfenati, 2011). We found that Syn is likely a direct target of Da since Da binds to *Syn* promoter, and both silencing and overexpression of *da* changes *Syn* levels.

We also sought out to rescue the learning phenotype caused by *da* silencing. For this we fed the larvae with resveratrol or SAHA. In luciferase reporter assay in primary neuronal cultures resveratrol improves transactivational capability of TCF4 significantly (unpublished data from our lab). Resveratrol inhibits cAMP-degrading phosphodiesterases which leads to elevated cAMP levels (Park et al., 2012) and TCF4-dependent transcription upon neuronal activity is activated by cAMP-PKA pathway mediated phosphorylation of TCF4 (Sepp et al., 2017). It is plausible that Da could also be regulated by phosphorylation by PKA and therefore resveratrol could improve Da transactivational capability. Indeed, resveratrol had a moderate positive effect on learning and memory of the *da* knockdown larvae. Whether this effect is through cAMP-PKA pathway is yet to be verified. SAHA is a histone deacetylase inhibitor which improves learning and memory in *TCF4*(+/−) mice through the normalization of synaptic plasticity (Kennedy et al., 2016). Here we show that feeding SAHA to *Drosophila* larvae had also a moderate effect on learning.

We also made an attempt to rescue the impaired geotaxis of 30Y>Dcr2;da^RNAi^ flies using resveratrol or SAHA, but we could not get significant improvement. We administrated the drugs in the food substrate either during development or to adult flies. Getting resveratrol or SAHA during development in the larval stage and the amount ingested by the adults is probably not enough to rescue phenotypes caused by lowered levels of Da in the adults. If administrating the drug in the food substrate during development and to adults prior to testing is insufficient, other administration methods could be used like possibly starving the adult flies to increase the amounts of drug ingested (Pandey and Nichols, 2011). In a recent study in *Drosophila melanogaster* where genes associated with autism spectrum disorders and intellectual disability were suppressed, the knockdown of Da resulted in impaired habituation (Fenckova et al., 2018). Rescuing this habituation phenotype could be also tested for improvement with drugs.

In conclusion, this study demonstrates that the levels of TCF4 homolog Da are important for learning and memory of *Drosophila* larvae and that Da directly regulates the expression of Syn and dlg1. This gives novel insights into the mechanisms of PTHS and this learning deficit model can be used for screening therapeutics for PTHS.

## Materials and methods

### *Drosophila* stocks

All *Drosophila* stocks and crosses were kept on malt and semolina based food with 12 h light and dark daily rhythms at 25°C with 60% humidity. *Drosophila* strains used in this study were CantonS (a gift from dr. Bertram Gerber), 201Y-Gal4 (Bloomington Drosophila Stock Center (BDSC #4440) and 30Y-Gal4 (BDSC #30818) (Yao Yang et al., 1995) were gifts from Mark Fortini. GMR12B08-Gal4 (BDSC #48489) (Pfeiffer et al., 2008), *elav*-Gal4 (BDSC #8760) (Luo et al., 1994), *repo*-Gal4 (BDSC #7415) (Sepp et al., 2001), UAS-*Dcr2*;*Pin*^*1*^/CyO (BDSC #24644) (Dietzl et al., 2007), UAS-nlsGFP (BDSC #4776), UAS-*da*^*G*^ (BDSC #37291), *da* mutant line *da*^*10*^ (BDSC #5531) (Caudy et al., 1988) from Bloomington Stock Center at Indiana University, USA. The following transgenic lines were generated in this study: 3xFLAG-*da*^*2M4*^ and sfGFP-*da*^*4M1*^.

### Endogeneous tagging of Da by CRISPR/Cas9

The coding sequence for 3xFLAG- or sfGFP-tag was inserted to the 5’ coding region of *da* gene using CRISPR-Cas9 technology. Genomic sequence around the tag was following: 5’ ATGGCGA**CCA**GTG|ACGATGAGCC 3’ (PAM sequence shown as bold and the cut site marked with |). For the higher mutagenesis rate a specific fruit fly line for guide RNA production was created. Partially overlapping oligonucleotides 5’-CTTCGTGCATCGGCTCATCGTCAC-3’ and 5’-AAACTGGACGATGAGCCGATGCAC-3’, designed to target the N-terminus of Da protein, were cloned downstream the Polymerase III U6:2 promoter in the pCFD2-dU6:2gRNA plasmid (Addgene #49409). Transgenic flies expressing gRNA-s were created by injecting the generated plasmid into PBac{yellow^+^-attP-9A}VK00027 (BDSC #9744) fly strain embryos. For donor plasmid generation, pHD-3xFLAG-ScarlessDsRed (a gift from Kate O’Connor-Giles (DGRC)) or pHD-sfGFP-ScarlessDsRed (a gift from Kate O’Connor-Giles (DGRC)) and Gibson cloning was used. The following primer pairs were used for amplification of upstream and downstream homology arms: upst F5 CGGCCGCGAATTCGCCCTTGGTTGTGAATCAGGTGTAGAAACA and upst_R GCCGGAACCTCCAGATCCACCACTGGTCGCCATTTCAGCA, dwns_F TTCTGGTGGTTCAGGAGGTTACGATGAGCCGATGCACTTG and dwns_R GTTTAAACGAATTCGCCCTTAACGCCCTGGAACACCGAGG.

After verification, the obtained donor plasmids pHD-*da*-3xFLAG-ScarlessDsRed and pHD-*da*-sfGFP-ScarlessDsRed were injected into F_1_ embryos from a cross between *da*-gRNA (our gRNA expressing transgenic strain) and y^1^M{w^+mC^=nos-Cas9.P}ZH-2A w* (BDSC #54591) fly strains. All embryo injections were ordered from BestGene Inc. (USA).

dsRed cassette was removed from selected progeny by crossing to PB transposase line *Herm{3xP3-ECFP,αtub-piggyBacK10}M10* (BDSC #32073) (Horn et al., 2003). The obtained 3xFLAG*-da* and sfGFP*-da* lines were verified by sequencing.

### RNA isolation and cloning

RNA from *3xFLAG-da* or *sfGFP-da Drosophila* embryos was isolated using RNeasy Mini Kit (Qiagen) according to the manufacturer’s protocol. cDNA was synthesized using 2µg of RNA. Primer sequences for cloning were 5’ ACTAGTTGAAGTCGACTGGAC 3’ and 5’ CCAGGTCCTCCAATTCCACC 3’. PCR products containing either *3xFLAG-da* or *sfGFP-da* cDNA sequences were sequenced and cloned into pCDNA3.1 expression vector (Tamberg et al., 2015) using BcuI (SpeI, 10 U; Thermo Scientific) and BstII (Eco 91I, 10 U; Thermo Scientific) restriction enzymes. The pcDNA3.1 constructs encoding Da, and reporter vectors pGL4.29[luc2P/12µE5/Hygro], pGL4[hRlucP/PGK], pGL4.83[hRlucP/EF1α/Puro] have been described previously (Sepp et al., 2011; Sepp et al., 2012; Tamberg et al., 2015).

### Cell culture and transfections and luciferase reporter assay

Human embryonic kidney cells HEK-293 obtained from ATCC (LGC Standards GmbH, Wesel, Germany) and routinely tested for contamination were grown in MEM (Capricorn Scientific) supplemented with 10% fetal bovine serum (PAA Laboratories), 100 U/ml penicillin, and 0.1 mg/ml streptomycin (Gibco). For transfection 0.375 µg of DNA and 0.75 µg of PEI (Sigma-Aldrich) were used per well of a 48-well plate or scaled up accordingly. For cotransfections, equal amounts of pGL4.29[luc2P/12µE5/Hygro], pGL4[hRlucP/min/Hygro], and effector constructs were used. Luciferase assays were performed as described previously (Sepp et al., 2011) using Passive Lysis Buffer (Promega) and the Dual-Glo Luciferase Assay (Promega). Cells were lysed at 24 h after transfection. For data analysis, background signals from untransfected cells were subtracted and firefly luciferase signals were normalised to *Renilla* luciferase signals. The data was then log-transformed, auto-scaled, means and standard deviations were calculated and Student t-tests were performed. The data was back-transformed for graphical representation.

### Protein electrophoresis and Western blotting

For SDS-PAGE embryos, larvae, pupae, adult heads or larval brains were lysed in 2x SDS sample buffer. Equal amounts of protein were loaded to gel.

The following mouse monoclonal antibodies were obtained from the Developmental Studies Hybridoma Bank (created by the NICHD of the NIH and maintained at The University of Iowa, Department of Biology, Iowa City, IA 52242): β-tubulin E7 (developed by Klymkowsky, M., dilution 1:3000); Synapsin SYNORF1 3C11 (developed by Buchner, E., 1:1000); discs large 1 4F3 (developed by Goodman, C., 1:2000), elav 9F8A9 (developed by Rubin, Gerald M., 1:1000), Bruchpilot nc82 (developed by Buchner, E., 1:100).

Other antibodies and dilutions used: mouse anti-Da dam109-10 1:10 (a gift from C. Cronmiller), mouse anti-FLAG M2 HRP-conjugated 1:6000 (Sigma-Aldrich), goat anti-mouse IgM HRP-conjugated secondary antibody (MilliporeSigma).

### Immunohistochemical staining

The anterior parts of 3rd instar larvae were dissected in PBS and fixed using 4% paraformaldehyde in PBS. Adult flies were first fixed in 4% paraformaldehyde in PBS and then dissected. Primary antibody labelling was performed over 72 hours on overhead rotator at 4°C in PBS with 0.5% TritonX-100. Used antibodies were as follows: mouse anti-FLAG M2 1:1000 (Sigma-Aldrich) and goat anti-mouse Alexa594 1:1000 (ImmunoResearch Laboratories). Secondary antibodies were preadsorbed to wt tissues before use. Incubation with secondary antibodies was performed for 3 h on overhead rotator at room temperature in PBS with 0.1% TritonX-100. The labelled larval brains were dissected and mounted in Vectashield mounting medium (Vector Laboratories). For image collection, Zeiss LSM 510 Meta confocal microscope with Pln Apo 20×/0.8 DICII objective (Carl Zeiss Microscopy) was used. Suitable layers were selected using Imaris (Bitplane Inc.) software.

### Appetitive associative learning assay

Appetitive associative memory assay in the *Drosophila* larvae was performed as previously described (Michels et al., 2017). Shortly, the larvae were trained three times for 5 minutes on Petri dishes, where one odor - amyl acetate (AM) was presented with plain agar and the other odor – octanol (OCT) with agar containing fructose as a reward. Then the larvae were placed in the midline of a plain agar plate and given a choice between the two odors placed on separate halves of the Petri dish and after three minutes larvae were counted on each half of the Petri dish. Then reciprocal training was performed with AM and fructose and OCT with plain agar. Using data from two reciprocally trained tests the performance index (PI) was calculated PI = (PREF AM_AM+/OCT_ - PREF AM_AM/OCT+_) / 2. The odors and the reward were presented in four different orders to eliminate any non-specific preferences. All together 12 training and test cycles were conducted per genotype, each time with new larvae, PI-s were calculated and used for statistical analysis. The PI-s were visualized as box-whisker plots, which show the median, the 10% - 90% quantiles, and the 25% - 75% quantiles. For statistical analysis inside one genotype a one-sample sign test was applied with an error threshold of smaller than 5%. For pairwise U-tests Bonferroni correction was used. SAHA was dissolved in dimethyl sulfoxide (DMSO) and same concentration of DMSO (0.1%) was used in the food substrate for a control. Resveratrol was dissolved in 96% ethanol and 1% ethanol in the food was used for the control.

### Chromatin immunoprecipitation

Chromatin preparations were carried out as previously described (Chanas et al., 2004). Approximately 150 mg of adult heads were collected on dry ice and homogenized in buffer A1 (60 mM KCl, 15 mM NaCl, 4 mM MgCl_2_, 15 mM HEPES pH 7.6, 0.5% Triton X-100, 0.5 mM DTT, 10 mM sodium butyrate, 1 × EDTA-free protease inhibitor cocktail (Roche)) + 1.8% formaldehyde in room temperature using first KONTES pellet pestle followed by three strokes using Dounce homogenizer with a loose pestle. Homogenate was incubated 15 minutes and glycin was added to 225 mM followed by 5 minutes incubation. Homogenate was then centrifuged 5 minutes at 4000 g at 4°C and supernatant discarded. Pellet was washed three times with 3 ml A1 followed by a wash with 3 ml of lysis buffer (14 mM NaCl, 15 mM HEPES pH 7.6, 1 mM EDTA, 0.5 mM EGTA, 1% Trition X-100, 0.5 mM DTT, 0.1% sodium deoxycholate, 0.05% SDS, 10 mM sodium butyrate, 1 × EDTA-free protease inhibitor cocktail (Roche)). Cross-linked material was resuspended in 0.5 ml of lysis buffer + 0.1% SDS and 0.5% N-lauroylsarcosine and incubated 10 minutes at 4°C on a rotator followed by sonication using SONICS VibraCell on 70% amplitude 15 seconds intervals 30 times. Cross-linked material was then rotated 10 minutes at 4°C and centrifuged 5 minutes at room temperature at maximum speed. Supernatant was transferred to a new tube and 0.5 ml of lysis buffer was added to the pellet followed by rotation and centrifugation. Supernatants were combined and centrifuged 2×10 minutes at maximum speed. Chromatin extract was transferred to Microcon DNA Fast Flow Centrifugal Filter Units (Merck Millipore) blocked with 1mg/ml BSA in PBS and purified using lysis buffer. The volume of chromatin extract was brought to 1 ml using lysis buffer. Protein concentrations were determined using BCA assay (Pierce).

After removing equal amounts of inputs chromatin extracts were diluted 10x using dilution buffer (1% Triton X-100, 150 mM NaCl, 2 mM EDTA pH 8.0, 20 mM Tris-HCl pH 8.0, 1 × EDTA-free protease inhibitor cocktail (Roche)) and added to 50 µl of Dynabeads Protein G (Invitrogen) beads that were previously incubated with 5 µg of monoclonal anti-FLAG M2 antibody (Sigma-Aldrich) in 0.05% PBS Tween20 overnight. Chromatin immunoprecipitation (ChIP) was carried out overnight at 4°C. Beads with chromatin were then washed in wash buffer (1% Triton X-100, 0.1% SDS, 150 mM NaCl, 2 mM EDTA pH 8.0, 20 mM Tris-HCl pH 8.0, 1 × EDTA-free protease inhibitor cocktail (Roche)) using a magnetic rack for 10 minutes three times at 4°C on a rotator followed by final wash buffer (1% Triton X-100, 0.1% SDS, 500 mM NaCl, 2 mM EDTA pH 8.0, 20 mM Tris-HCl pH 8.0, 1 × EDTA-free protease inhibitor cocktail (Roche)). Chromatin was eluted using 3×50 µl of elution buffer (1% SDS, 100 mM NaHCO_3_, 1 mM EDTA) 3×10 minutes at 37°C. The volume of inputs was brought to 150 µl with elution buffer. For decrosslinking 8 µl of 5 M NaCl was added and the samples were incubated at 65°C overnight. Then 2 µl of RNase A (10 mg/ml) was added and incubated at 37°C for 30 minutes followed by incubation with 2 µl of EDTA (0.5M) and 4 µl Proteinase K (10 mg/ml) at 45°C for 30 minutes. DNA was extracted using QIAquick PCR Purification Kit (Qiagen).

### Quantitative PCR

For RT-qPCR 15 heads were collected from 2-3 days old adult flies on dry ice. RNA was extracted using RNeasy Mini Kit (Qiagen). cDNA was synthesized with Superscript IV reverse transcriptase (Invitrogen) and oligo(dT)_20_ primers. Quantitative PCR was performed using LightCycler 480 II (Roche) with Hot FIREPol EvaGreen qPCR Mix Plus (Solis Biodyne). Primer sequences are shown in Supplementary table 1.

### Negative geotaxis assay

10 females and males were separated to fresh vials 48 hours before the assay to allow recovering from anesthesia. Prior to the test males and females from control and *da* silencing group were transferred to empty vials without anesthesia and closed with another upside down vial using sticky tape. The flies were knocked down three times on the table and a photo was taken after 10 seconds. The height of the vial was divided into 10 equal parts and the number of flies in each compartment was counted and average height was calculated. The experiment was repeated five times, each time with new flies. Average climbing heights were visualized using box-whisker plots, which show the median, the 10% - 90% quantiles, and the 25% - 75% quantiles. For statistical significance pairwise U-tests were used.

## Acknowledgements

We thank Mark Fortini, Bertram Gerber and Claire Cronmiller for sharing fly stocks and reagents. We thank Epp Väli and Jan Erik Alliksaar for technical assistance. We are grateful to Jürgen Tuvikene, Allan-Hermann Pool, and Richard Tamme for critical comments on manuscript. Stocks obtained from the Bloomington Drosophila Stock Center (NIH P40OD018537) were used in this study. We acknowledge the Drosophila Genomics Resource Center supported by NIH grant 2P40OD010949.

## Competing interests

No competing interests declared.

## Funding

This study was supported by Estonian Research Council (institutional research funding IUT19-18), European Union through the European Regional Development Fund (Project No. 2014-2020.4.01.15-0012) and H2020-MSCA-RISE-2016 (EU734791), Pitt Hopkins Research Foundation and Million Dollar Bike Ride Pilot Grant Program for Rare Disease Research at UPenn Orphan Disease Center (grants MDBR-16-122-PHP and MDBR-17-127-Pitt Hopkins).

## Author contributions statement

Conceptualization: L.T., M.P.; Methodology: L.T., M.J., M.P.; Formal analysis: L.T., A.Sh., C.S.K.; Investigation: L.T., M.J., K.S., A.Sh., C.S.K., M.P.; Resources: T.T.; Writing – original draft preparation: L.T.; Writing – review and editing: L.T., A.S., T.T., M.P.; Visualization: L.T., M.P.; Supervision: L.T., A.S., M.P.; Project administration: T.T., M.P.; Funding acquisition: T.T;

## Notes

#### Summary of Updates

The Supplemental figures and table was added

## References

Amiel, J., Rio, M., de Pontual, L., Redon, R., Malan, V., Boddaert, N., Plouin, P., Carter, N. P., Lyonnet, S., Munnich, A., et al. (2007). Mutations in TCF4, encoding a class I basic helix-loop-helix transcription factor, are responsible for Pitt-Hopkins syndrome, a severe epileptic encephalopathy associated with autonomic dysfunction. Am. J. Hum. Genet. 80, 988–993.

Andrade-Zapata, I. and Baonza, A. (2014). The bHLH factors extramacrochaetae and daughterless control cell cycle in Drosophila imaginal discs through the transcriptional regulation of the Cdc25 phosphatase string. PLoS Genet. 10, e1004233.

Aso, Y., Grübel, K., Busch, S., Friedrich, A. B., Siwanowicz, I. and Tanimoto, H. (2009). The mushroom body of adult Drosophila characterized by GAL4 drivers. J. Neurogenet. 23, 156–172.

Bardin, A. J., Perdigoto, C. N., Southall, T. D., Brand, A. H. and Schweisguth, F. (2010). Transcriptional control of stem cell maintenance in the Drosophila intestine. Dev. Camb. Engl. 137, 705–714.

Benfenati, F. (2011). Synapsins—Molecular function, development and disease. Semin. Cell Dev. Biol. 22, 377.

Bhattacharya, A. and Baker, N. E. (2011). A network of broadly expressed HLH genes regulates tissue-specific cell fates. Cell 147, 881–892.

Bhattacharya, A. and Baker, N. E. (2012). The role of the bHLH protein hairy in morphogenetic furrow progression in the developing Drosophila eye. PloS One 7, e47503.

Brand, A. H. and Perrimon, N. (1993). Targeted gene expression as a means of altering cell fates and generating dominant phenotypes. Dev. Camb. Engl. 118, 401–415.

Brockschmidt, A., Todt, U., Ryu, S., Hoischen, A., Landwehr, C., Birnbaum, S., Frenck, W., Radlwimmer, B., Lichter, P., Engels, H., et al. (2007). Severe mental retardation with breathing abnormalities (Pitt-Hopkins syndrome) is caused by haploinsufficiency of the neuronal bHLH transcription factor TCF4. Hum. Mol. Genet. 16, 1488–1494.

Brown, N. L., Paddock, S. W., Sattler, C. A., Cronmiller, C., Thomas, B. J. and Carroll, S. B. (1996). daughterless is required for Drosophila photoreceptor cell determination, eye morphogenesis, and cell cycle progression. Dev. Biol. 179, 65–78.

Cabrera, C. V. and Alonso, M. C. (1991). Transcriptional activation by heterodimers of the achaete-scute and daughterless gene products of Drosophila. EMBO J. 10, 2965–2973.

Castanon, I., Von Stetina, S., Kass, J. and Baylies, M. K. (2001). Dimerization partners determine the activity of the Twist bHLH protein during Drosophila mesoderm development. Dev. Camb. Engl. 128, 3145–3159.

Caudy, M., Vässin, H., Brand, M., Tuma, R., Jan, L. Y. and Jan, Y. N. (1988). daughterless, a Drosophila gene essential for both neurogenesis and sex determination, has sequence similarities to myc and the achaete-scute complex. Cell 55, 1061–1067.

Chanas, G., Lavrov, S., Iral, F., Cavalli, G. and Maschat, F. (2004). Engrailed and polyhomeotic maintain posterior cell identity through cubitus-interruptus regulation. Dev. Biol. 272, 522–535.

Chen, T., Wu, Q., Zhang, Y., Lu, T., Yue, W. and Zhang, D. (2016). Tcf4 Controls Neuronal Migration of the Cerebral Cortex through Regulation of Bmp7. Front. Mol. Neurosci. 9, 94.

Cisse, B., Caton, M. L., Lehner, M., Maeda, T., Scheu, S., Locksley, R., Holmberg, D., Zweier, C., den Hollander, N. S., Kant, S. G., et al. (2008). Transcription factor E2-2 is an essential and specific regulator of plasmacytoid dendritic cell development. Cell 135, 37–48.

Cline, T. W. (1978). Two closely linked mutations in Drosophila melanogaster that are lethal to opposite sexes and interact with daughterless. Genetics 90, 683–698.

Cronmiller, C. and Cummings, C. A. (1993). The daughterless gene product in Drosophila is a nuclear protein that is broadly expressed throughout the organism during development. Mech. Dev. 42, 159–169.

Crux, S., Herms, J. and Dorostkar, M. M. (2018). Tcf4 regulates dendritic spine density and morphology in the adult brain. PLoS ONE 13,.

Cummings, C. A. and Cronmiller, C. (1994). The daughterless gene functions together with Notch and Delta in the control of ovarian follicle development in Drosophila. Dev. Camb. Engl. 120, 381–394.

Cuthbert, P. C., Stanford, L. E., Coba, M. P., Ainge, J. A., Fink, A. E., Opazo, P., Delgado, J. Y., Komiyama, N. H., O’Dell, T. J. and Grant, S. G. N. (2007). Synapse-Associated Protein 102/dlgh3 Couples the NMDA Receptor to Specific Plasticity Pathways and Learning Strategies. J. Neurosci. Off. J. Soc. Neurosci. 27, 2673–2682.

Diegelmann, S., Klagges, B., Michels, B., Schleyer, M. and Gerber, B. (2013). Maggot learning and Synapsin function. J. Exp. Biol. 216, 939–951.

Dietzl, G., Chen, D., Schnorrer, F., Su, K.-C., Barinova, Y., Fellner, M., Gasser, B., Kinsey, K., Oppel, S., Scheiblauer, S., et al. (2007). A genome-wide transgenic RNAi library for conditional gene inactivation in *Drosophila*. Nature 448, 151–156.

Ekins, S., Gerlach, J., Zorn, K. M., Antonio, B. M., Lin, Z. and Gerlach, A. (2019). Repurposing Approved Drugs as Inhibitors of Kv7.1 and Nav1.8 to Treat Pitt Hopkins Syndrome. Pharm. Res. 36, 137.

Fenckova, M., Asztalos, L., Cizek, P., Singgih, E. L., Blok, L. E. R., Glennon, J. C., IntHout, J., Zweier, C., Eichler, E. E., Bernier, R., et al. (2018). Integrative Cross-species Analyses Suggest Deficits in Habituation Learning as a Widely Affected Mechanism in Intellectual Disability and Autism Spectrum Disorders. bioRxiv 285981.

Forrest, M. P., Waite, A. J., Martin-Rendon, E. and Blake, D. J. (2013). Knockdown of human TCF4 affects multiple signaling pathways involved in cell survival, epithelial to mesenchymal transition and neuronal differentiation. PloS One 8, e73169.

Forrest, M. P., Hill, M. J., Kavanagh, D. H., Tansey, K. E., Waite, A. J. and Blake, D. J. (2018). The Psychiatric Risk Gene Transcription Factor 4 (TCF4) Regulates Neurodevelopmental Pathways Associated With Schizophrenia, Autism, and Intellectual Disability. Schizophr. Bull. 44, 1100–1110.

Garcia, C. C., Blair, H. J., Seager, M., Coulthard, A., Tennant, S., Buddles, M., Curtis, A. and Goodship, J. A. (2004). Identification of a mutation in synapsin I, a synaptic vesicle protein, in a family with epilepsy. J. Med. Genet. 41, 183–186.

Giebel, B., Stüttem, I., Hinz, U. and Campos-Ortega, J. A. (1997). Lethal of Scute requires overexpression of Daughterless to elicit ectopic neuronal development during embryogenesis in Drosophila. Mech. Dev. 63, 75–87.

Gitler, D., Takagishi, Y., Feng, J., Ren, Y., Rodriguiz, R. M., Wetsel, W. C., Greengard, P. and Augustine, G. J. (2004). Different Presynaptic Roles of Synapsins at Excitatory and Inhibitory Synapses. J. Neurosci. 24, 11368–11380.

Grajkowska, L. T., Ceribelli, M., Lau, C. M., Warren, M. E., Tiniakou, I., Higa, S. N., Bunin, A., Haecker, H., Mirny, L. A., Staudt, L. M., et al. (2017). Isoform-specific expression and feedback regulation of E protein TCF4 control dendritic cell lineage specification. Immunity 46, 65–77.

Hennig, K. M., Fass, D. M., Zhao, W.-N., Sheridan, S. D., Fu, T., Erdin, S., Stortchevoi, A., Lucente, D., Cody, J. D., Sweetser, D., et al. (2017). WNT/β-Catenin Pathway and Epigenetic Mechanisms Regulate the Pitt-Hopkins Syndrome and Schizophrenia Risk Gene TCF4. Mol. Neuropsychiatry 3, 53–71.

Hill, M. J., Killick, R., Navarrete, K., Maruszak, A., McLaughlin, G. M., Williams, B. P. and Bray, N. J. (2017). Knockdown of the schizophrenia susceptibility gene TCF4 alters gene expression and proliferation of progenitor cells from the developing human neocortex. J. Psychiatry Neurosci. JPN 42, 181–188.

Horn, C., Offen, N., Nystedt, S., Häcker, U. and Wimmer, E. A. (2003). piggyBac-based insertional mutagenesis and enhancer detection as a tool for functional insect genomics. Genetics 163, 647–661.

Jung, M., Häberle, B. M., Tschaikowsky, T., Wittmann, M.-T., Balta, E.-A., Stadler, V.-C., Zweier, C., Dörfler, A., Gloeckner, C. J. and Lie, D. C. (2018). Analysis of the expression pattern of the schizophrenia-risk and intellectual disability gene TCF4 in the developing and adult brain suggests a role in development and plasticity of cortical and hippocampal neurons. Mol. Autism 9,.

Kennedy, A. J., Rahn, E. J., Paulukaitis, B. S., Savell, K. E., Kordasiewicz, H. B., Wang, J., Lewis, J. W., Posey, J., Strange, S. K., Guzman-Karlsson, M. C., et al. (2016). Tcf4 Regulates Synaptic Plasticity, DNA Methylation, and Memory Function. Cell Rep. 16, 2666–2685.

Kepa, A., Medina, L. M., Erk, S., Srivastava, D. P., Fernandes, A., Toro, R., Lévi, S., Ruggeri, B., Fernandes, C., Degenhardt, F., et al. (2017). Associations of the intellectual disability gene MYT1L with helix-loop-helix gene expression, hippocampus volume and hippocampus activation during memory retrieval. Neuropsychopharmacol. Off. Publ. Am. Coll. Neuropsychopharmacol. 42, 2516–2526.

King-Jones, K., Korge, G. and Lehmann, M. (1999). The helix-loop-helix proteins dAP-4 and daughterless bind both in vitro and in vivo to SEBP3 sites required for transcriptional activation of the Drosophila gene Sgs-4. J. Mol. Biol. 291, 71–82.

Klagges, B. R. E., Heimbeck, G., Godenschwege, T. A., Hofbauer, A., Pflugfelder, G. O., Reifegerste, R., Reisch, D., Schaupp, M., Buchner, S. and Buchner, E. (1996). Invertebrate Synapsins: A Single Gene Codes for Several Isoforms in Drosophila. J. Neurosci. 16, 3154–3165.

Lennertz, L., Rujescu, D., Wagner, M., Frommann, I., Schulze-Rauschenbach, S., Schuhmacher, A., Landsberg, M. W., Franke, P., Möller, H.-J., Wölwer, W., et al. (2011a). Novel schizophrenia risk gene TCF4 influences verbal learning and memory functioning in schizophrenia patients. Neuropsychobiology 63, 131–136.

Lennertz, L., Quednow, B. B., Benninghoff, J., Wagner, M., Maier, W. and Mössner, R. (2011b). Impact of TCF4 on the genetics of schizophrenia. Eur. Arch. Psychiatry Clin. Neurosci. 261 Suppl 2, S161–165.

Li, K. and Baker, N. E. (2018). Regulation of the Drosophila ID protein Extra macrochaetae by proneural dimerization partners. eLife.

Li, H., Zhu, Y., Morozov, Y. M., Chen, X., Page, S. C., Rannals, M. D., Maher, B. J. and Rakic, P. (2019). Disruption of TCF4 regulatory networks leads to abnormal cortical development and mental disabilities. Mol. Psychiatry.

Luo, L., Liao, Y. J., Jan, L. Y. and Jan, Y. N. (1994). Distinct morphogenetic functions of similar small GTPases: Drosophila Drac1 is involved in axonal outgrowth and myoblast fusion. Genes Dev. 8, 1787–1802.

MacArthur, S., Li, X.-Y., Li, J., Brown, J. B., Chu, H. C., Zeng, L., Grondona, B. P., Hechmer, A., Simirenko, L., Keränen, S. V., et al. (2009). Developmental roles of 21 Drosophila transcription factors are determined by quantitative differences in binding to an overlapping set of thousands of genomic regions. Genome Biol. 10, R80.

Massari, M. E. and Murre, C. (2000). Helix-Loop-Helix Proteins: Regulators of Transcription in Eucaryotic Organisms. Mol. Cell. Biol. 20, 429–440.

Michels, B., Diegelmann, S., Tanimoto, H., Schwenkert, I., Buchner, E. and Gerber, B. (2005). A role for Synapsin in associative learning: The Drosophila larva as a study case. Learn. Mem. 12, 224–231.

Michels, B., Chen, Y., Saumweber, T., Mishra, D., Tanimoto, H., Schmid, B., Engmann, O. and Gerber, B. (2011). Cellular site and molecular mode of synapsin action in associative learning. Learn. Mem. 18, 332–344.

Michels, B., Saumweber, T., Biernacki, R., Thum, J., Glasgow, R. D. V., Schleyer, M., Chen, Y., Eschbach, C., Stocker, R. F., Toshima, N., et al. (2017). Pavlovian Conditioning of Larval Drosophila: An Illustrated, Multilingual, Hands-On Manual for Odor-Taste Associative Learning in Maggots. Front. Behav. Neurosci. 11,.

Murre, C., Bain, G., van Dijk, M. A., Engel, I., Furnari, B. A., Massari, M. E., Matthews, J. R., Quong, M. W., Rivera, R. R. and Stuiver, M. H. (1994). Structure and function of helix-loop-helix proteins. Biochim. Biophys. Acta 1218, 129–135.

Page, S. C., Hamersky, G. R., Gallo, R. A., Rannals, M. D., Calcaterra, N. E., Campbell, M. N., Mayfield, B., Briley, A., Phan, B. N., Jaffe, A. E., et al. (2018). The schizophrenia- and autism-associated gene, transcription factor 4 regulates the columnar distribution of layer 2/3 prefrontal pyramidal neurons in an activity-dependent manner. Mol. Psychiatry 23, 304–315.

Pandey, U. B. and Nichols, C. D. (2011). Human Disease Models in Drosophila melanogaster and the Role of the Fly in Therapeutic Drug Discovery. Pharmacol. Rev. 63, 411–436.

Park, D., Shafer, O. T., Shepherd, S. P., Suh, H., Trigg, J. S. and Taghert, P. H. (2008). The Drosophila basic helix-loop-helix protein DIMMED directly activates PHM, a gene encoding a neuropeptide-amidating enzyme. Mol. Cell. Biol. 28, 410–421.

Park, S.-J., Ahmad, F., Philp, A., Baar, K., Williams, T., Luo, H., Ke, H., Rehmann, H., Taussig, R., Brown, A. L., et al. (2012). Resveratrol ameliorates aging-related metabolic phenotypes by inhibiting cAMP phosphodiesterases. Cell 148, 421–433.

Pfeiffer, B. D., Jenett, A., Hammonds, A. S., Ngo, T.-T. B., Misra, S., Murphy, C., Scully, A., Carlson, J. W., Wan, K. H., Laverty, T. R., et al. (2008). Tools for neuroanatomy and neurogenetics in Drosophila. Proc. Natl. Acad. Sci. U. S. A. 105, 9715–9720.

Powell, L. M., Deaton, A. M., Wear, M. A. and Jarman, A. P. (2008). Specificity of Atonal and Scute bHLH factors: analysis of cognate E box binding sites and the influence of Senseless. Genes Cells Devoted Mol. Cell. Mech. 13, 915–929.

Quednow, B. B., Ettinger, U., Mössner, R., Rujescu, D., Giegling, I., Collier, D. A., Schmechtig, A., Kühn, K.-U., Möller, H.-J., Maier, W., et al. (2011). The schizophrenia risk allele C of the TCF4 rs9960767 polymorphism disrupts sensorimotor gating in schizophrenia spectrum and healthy volunteers. J. Neurosci. Off. J. Soc. Neurosci. 31, 6684–6691.

Rannals, M. D. and Maher, B. J. (2017). Molecular Mechanisms of Transcription Factor 4 in Pitt Hopkins Syndrome. Curr. Genet. Med. Rep. 5, 1–7.

Rannals, M. D., Page, S. C., Campbell, M. N., Gallo, R. A., Mayfield, B. and Maher, B. J. (2016). Neurodevelopmental models of transcription factor 4 deficiency converge on a common ion channel as a potential therapeutic target for Pitt Hopkins syndrome. Rare Dis. Austin Tex 4, e1220468.

Sepp, K. J., Schulte, J. and Auld, V. J. (2001). Peripheral glia direct axon guidance across the CNS/PNS transition zone. Dev. Biol. 238, 47–63.

Sepp, M., Kannike, K., Eesmaa, A., Urb, M. and Timmusk, T. (2011). Functional diversity of human basic helix-loop-helix transcription factor TCF4 isoforms generated by alternative 5’ exon usage and splicing. PloS One 6, e22138.

Sepp, M., Pruunsild, P. and Timmusk, T. (2012). Pitt-Hopkins syndrome-associated mutations in TCF4 lead to variable impairment of the transcription factor function ranging from hypomorphic to dominant-negative effects. Hum. Mol. Genet. 21, 2873–2888.

Sepp, M., Vihma, H., Nurm, K., Urb, M., Page, S. C., Roots, K., Hark, A., Maher, B. J., Pruunsild, P. and Timmusk, T. (2017). The Intellectual Disability and Schizophrenia Associated Transcription Factor TCF4 Is Regulated by Neuronal Activity and Protein Kinase A. J. Neurosci. 37, 10516–10527.

Silva, A. J., Rosahl, T. W., Chapman, P. F., Marowitz, Z., Friedman, E., Frankland, P. W., Cestari, V., Cioffi, D., Südhof, T. C. and Bourtchuladze, R. (1996). Impaired learning in mice with abnormal short-lived plasticity. Curr. Biol. CB 6, 1509–1518.

Smith, J. E. and Cronmiller, C. (2001). The Drosophila daughterless gene autoregulates and is controlled by both positive and negative cis regulation. Dev. Camb. Engl. 128, 4705–4714.

Smith, J. E., Cummings, C. A. and Cronmiller, C. (2002). Daughterless coordinates somatic cell proliferation, differentiation and germline cyst survival during follicle formation in Drosophila. Dev. Camb. Engl. 129, 3255–3267.

Sun, J., Xu, A. Q., Giraud, J., Poppinga, H., Riemensperger, T., Fiala, A. and Birman, S. (2018). Neural Control of Startle-Induced Locomotion by the Mushroom Bodies and Associated Neurons in Drosophila. Front. Syst. Neurosci. 12,.

Talkowski, M. E., Rosenfeld, J. A., Blumenthal, I., Pillalamarri, V., Chiang, C., Heilbut, A., Ernst, C., Hanscom, C., Rossin, E., Lindgren, A. M., et al. (2012). Sequencing chromosomal abnormalities reveals neurodevelopmental loci that confer risk across diagnostic boundaries. Cell 149, 525–537.

Tamberg, L., Sepp, M., Timmusk, T. and Palgi, M. (2015). Introducing Pitt-Hopkins syndrome-associated mutations of TCF4 to Drosophila daughterless. Biol. Open 4, 1762–1771.

Tanaka-Matakatsu, M., Miller, J., Borger, D., Tang, W.-J. and Du, W. (2014). Daughterless homodimer synergizes with Eyeless to induce Atonal expression and retinal neuron differentiation. Dev. Biol. 392, 256–265.

Tarpey, P., Parnau, J., Blow, M., Woffendin, H., Bignell, G., Cox, C., Cox, J., Davies, H., Edkins, S., Holden, S., et al. (2004). Mutations in the DLG3 Gene Cause Nonsyndromic X-Linked Mental Retardation. Am. J. Hum. Genet. 75, 318–324.

Thaxton, C., Kloth, A. D., Clark, E. P., Moy, S. S., Chitwood, R. A. and Philpot, B. D. (2018). Common Pathophysiology in Multiple Mouse Models of Pitt–Hopkins Syndrome. J. Neurosci. 38, 918–936.

Wong, M.-C., Castanon, I. and Baylies, M. K. (2008). Daughterless dictates Twist activity in a context-dependent manner during somatic myogenesis. Dev. Biol. 317, 417–429.

Wu, Y., Bolduc, F. V., Bell, K., Tully, T., Fang, Y., Sehgal, A. and Fischer, J. A. (2008). A Drosophila model for Angelman syndrome. Proc. Natl. Acad. Sci. U. S. A. 105, 12399–12404.

Xia, H., Jahr, F. M., Kim, N.-K., Xie, L., Shabalin, A. A., Bryois, J., Sweet, D. H., Kronfol, M. M., Palasuberniam, P., McRae, M., et al. (2018). Building a schizophrenia genetic network: transcription factor 4 regulates genes involved in neuronal development and schizophrenia risk. Hum. Mol. Genet. 27, 3246–3256.

Yao Yang, M., Armstrong, J. D., Vilinsky, I., Strausfeld, N. J. and Kaiser, K. (1995). Subdivision of the drosophila mushroom bodies by enhancer-trap expression patterns. Neuron 15, 45–54.

Yasugi, T., Fischer, A., Jiang, Y., Reichert, H. and Knoblich, J. A. (2014). A Regulatory Transcriptional Loop Controls Proliferation and Differentiation in Drosophila Neural Stem Cells. PLoS ONE 9,.

Zanni, G., van Esch, H., Bensalem, A., Saillour, Y., Poirier, K., Castelnau, L., Ropers, H. H., de Brouwer, A. P. M., Laumonnier, F., Fryns, J.-P., et al. (2010). A novel mutation in the DLG3 gene encoding the synapse-associated protein 102 (SAP102) causes non-syndromic mental retardation. neurogenetics 11, 251–255.

Zollino, M., Zweier, C., Balkom, I. D. V., Sweetser, D. A., Alaimo, J., Bijlsma, E. K., Cody, J., Elsea, S. H., Giurgea, I., Macchiaiolo, M., et al. (2019). Diagnosis and management in Pitt-Hopkins syndrome: First international consensus statement. Clin. Genet. 95, 462–478.

Zweier, C., Peippo, M. M., Hoyer, J., Sousa, S., Bottani, A., Clayton-Smith, J., Reardon, W., Saraiva, J., Cabral, A., Gohring, I., et al. (2007). Haploinsufficiency of TCF4 causes syndromal mental retardation with intermittent hyperventilation (Pitt-Hopkins syndrome). Am. J. Hum. Genet. 80, 994–1001.

