## Supplementary material for "Daughterless, the *Drosophila* orthologue of TCF4, is required for associative learning and maintenance of synaptic proteome": Supllemental Figures 1, 2 and supplemental table 1

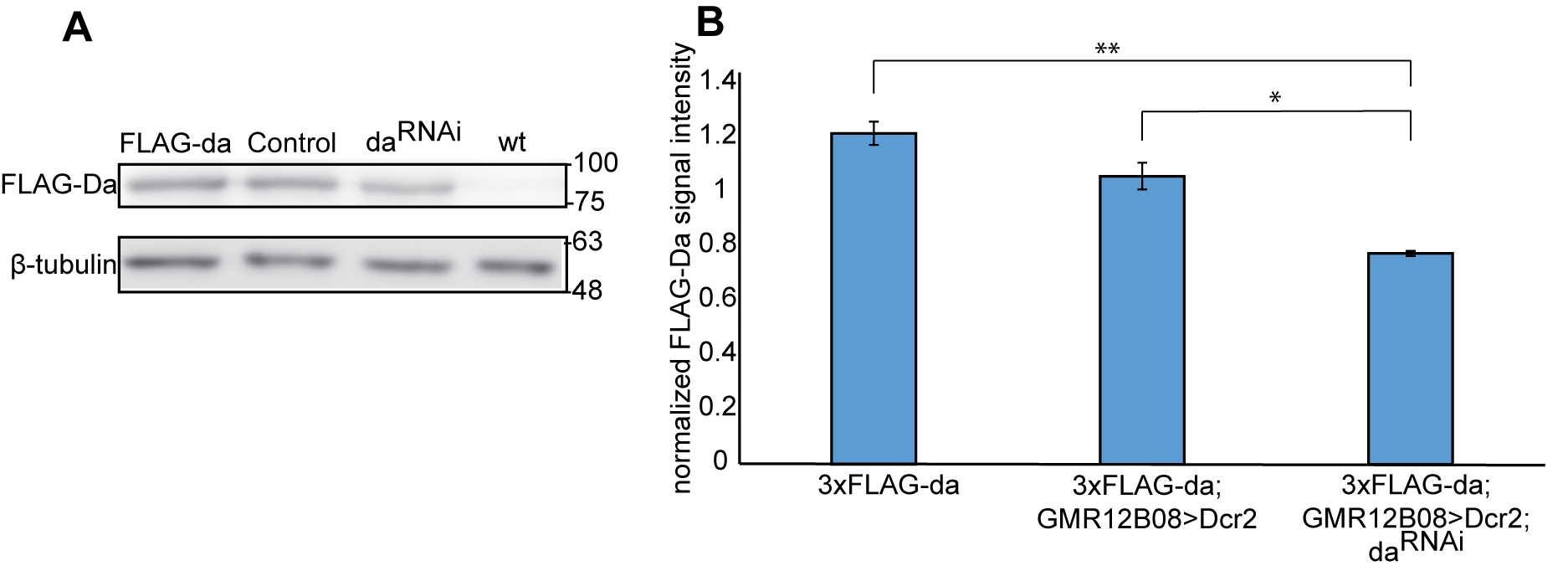

**Supplementary figure 1. Silencing of *da* in larval brain reduces Da levels**. A - Western blot analysis with anti-FLAG antibody of dissected third instar larval brains. Numbers on the right indicate molecular weight in kDa, 3xFLAG-da - larval brains where Da is tagged with 3xFLAG epitope, Control *– 3xFLAG*-*da*;GMR12B08>*Dcr2* larval brains, da^RNAi^ – *3xFLAG*-*da*;GMR12B08>*Dcr2*,*da^RNAi^* larval brains, wt - *white^1118^* larval brains for control lacking the 3xFLAG tag. B - upon silencing of *da* the Da expression levels were reduced about 35% compared to *3xFLAG*-*da* and about 25% compared to *3xFLAG*-*da*;GMR12B08>*Dcr2* larval brains. Shown are the results of densitometric analysis of Western blot, 3xFLAG-Da signals were normalized using β-tubulin signals. The mean results from three independent Western blots are shown. Error bars show standard errors. Statistical significance is shown with asterisks between the groups connected with lines. *P<0.05, **P<0.01, ***P<0.001.

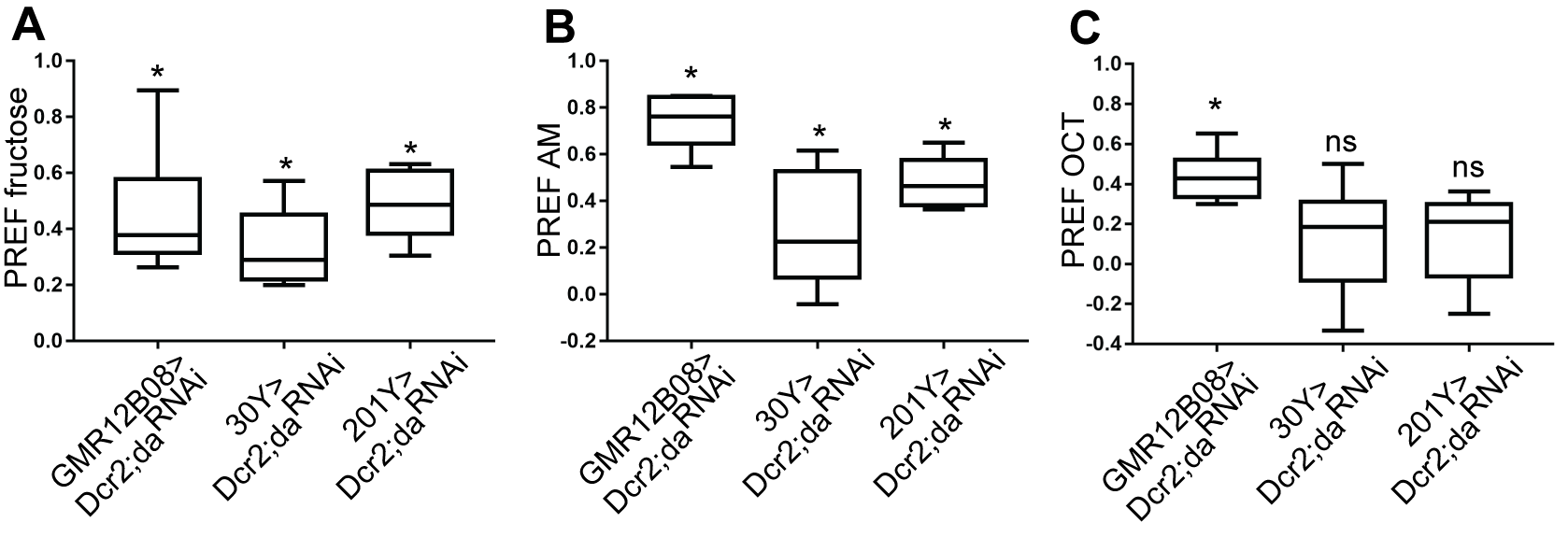

**Supplementary figure 2. Lowered levels of Da in the nervous system do not affect taste and smell sensing.** Larvae where *da* is silenced using GMR12B08-Gal4, 30Y-Gal4 or 201Y-Gal4 show preference towards fructose and amyl-acetate (AM) – A and B. Larvae do not move towards octanol (OCT) in the case of 30Y-Gal4 and 201Y-Gal4 (C). For statistical analysis one-sample sign test was used.

**Supplementary table 1. Used qPCR primers**

| **Primers** | **Forward** | **Reverse** |
| --- | --- | --- |
| **Ac (ChIP)** (Andrade-Zapata and Baonza, 2014) | ggtatcagggcctagggatcc | gatccttcagtgatgatgctgttg |
| **PHM (ChIP)** | cagaccgtaagtccaggttcc | cgggaaaaaccagagcctat |
| **SynI (ChIP)** | ttcgagtggaaatttgtgagc | gccgtgtttcgatttttggg |
| **SynII (ChIP)** | ccggctaatgacaccaggaa | ggattggcaggtagacgagg |
| **Dlg1I (ChIP)** | gctgcaattgcgaagctaca | tggttcgcaaggtgcgatag |
| **Dlg1II (ChIP)** | cgtcgccagatacacgagtt | cgtggtccagcttggtactc |
| **Dlg1III (ChIP)** | tcactgtattcggttcttgcct | ccagtctgtgtgagttggct |
| **Dlg1IV (ChIP)** | gctgggcatctgcgttctat | acttggctagtgatcctgctc |
| **DaQ** | ggtggctcaacgtcaacact | atcgtcactggtcgccattt |
| **AcQ** | aagcaaggagcatcgtcaca | agagtgattttcgctgccca |
| **PHMQ** | cgatctgtacttgtgcacgc | tatggtgggccgtgttcatc |
| **SynQ** | gcgagggtctgaacaatcca | gtggtcttgctgtctccgaa |
| **Dlg1Q1** | ccaagttgatggacggcgga | aggttcttctcgctcccgtt |
| **Dlg1Q2** | gcttgtttcaagcgctgtt | atttgcaaggtctccgctgt |
